# In-depth immunophenotyping with mass cytometry during TB treatment reveals non-canonical T-cell subsets associated with sputum culture conversion

**DOI:** 10.1101/2021.10.27.466125

**Authors:** Carole Chedid, Thibault Andrieu, Eka Kokhreidze, Nestani Tukvadze, Samanta Biswas, Md. Fahim Ather, Mohammad Khaja Mafij Uddin, Sayera Banu, Flavio De Maio, Giovanni Delogu, Hubert Endtz, Delia Goletti, Marc Vocanson, Oana Dumitrescu, Jonathan Hoffmann, Florence Ader

## Abstract

Tuberculosis (TB) is a difficult-to-treat infection because of multidrug regimen requirements based on drug susceptibility profiles and treatment observance issues. TB cure is defined by mycobacterial sterilization, technically complex to systematically assess. We hypothesized that microbiological outcome was associated with stage-specific immune changes in peripheral whole blood during TB treatment. The T-cell phenotypes of treated TB patients were prospectively characterized in a blinded fashion using mass cytometry after *Mycobacterium tuberculosis* (*Mtb*) antigen stimulation, and then correlated to sputum culture status. At two months of treatment, cytotoxic and terminally differentiated CD8^+^ T-cells were under-represented and naïve CD4^+^ T-cells were over-represented in positive-*versus* negative-sputum culture patients, regardless of *Mtb* drug susceptibility. At treatment completion, an antigen-driven T-cell immune shift towards differentiated subpopulations was associated with TB cure. Overall, we identified specific T-cell profiles associated with slow sputum converters, which brings new insights in TB prognostic biomarker research designed for clinical application.

**Summary:** In patients treated for pulmonary TB, high-dimensional immune profiling with mass cytometry revealed that *Mycobacterium tuberculosis* culture conversion is associated with newly characterized peripheral CD8^+^ T-cell phenotypes. This paves the way for new immune biomarkers associated with mycobacterial sterilization.

## Introduction

Tuberculosis (TB) is a leading cause of death of infectious origin, responsible for 1.5 million deaths worldwide in 2020 (World Health Organization Geneva, 2020). TB treatment regimens have toxic side effects (World Health Organization Geneva, 2019) requiring monitoring throughout treatment to adapt it and assess effectiveness. Pulmonary TB treatment monitoring relies on *Mycobacterium tuberculosis* (*Mtb*) detection in sputum samples (World Health Organization Geneva, 2018), which can be difficult to collect in later stages of treatment (Singhania et al., 2018). Smear microscopy yields highly sample- and operator-dependent results and has poor sensitivity (Parrish and Carroll, 2011). Sputum culture is the gold standard, although slow and requiring biosafety laboratory environments (Horne et al., 2010). Simultaneously, one of the main stakes in improving TB management is shortening TB treatment (Lienhardt et al., 2016). Overall, there is a need for novel non-sputum-based tools to monitor disease resolution and assess cure while remaining feasible in primary care settings (Goletti et al., 2018). Blood-based host immune biomarkers have recently gained interest in TB research as immune cells undergo phenotypic changes throughout the disease. Numerous past investigations have pointed to variations in the abundance and marker expression of several targeted subpopulations (Ahmed et al., 2018b; Adekambi et al., 2015a; Goletti et al., 2006b; Agrawal et al., 2018), in particular T-cells, which are pivotal effectors for *Mtb* clearance (Riou et al., 2020). However, this has been explored mostly in low-TB prevalence settings or with conventional flow cytometry, targeting a limited number of cell markers (Chiacchio et al., 2017; Musvosvi et al., 2018).

High-dimensional single-cell technologies such as mass cytometry enable the detection and quantification of a high number of cell markers (Gossez et al., 2018). This technique bypasses the limitations of spectral overlap by using monoclonal antibodies coupled to metal polymers, and has allowed high-dimensional exploration of the immune landscape in several domains (Kourelis et al., 2019; Rubin et al., 2019). It has been applied to immune profiling during TB treatment in a 2018 study by Roy Chowdhury and colleagues (Roy Chowdhury et al., 2018), in which the authors have provided a general overview of changes in the main immune blood cells during treatment.

Here, in a prospective, international cohort study of adult patients treated for pulmonary TB in high prevalence countries, peripheral blood T-cell immune-profiles were characterized using a 29-marker mass cytometry panel. In-depth T-cell phenotypical analysis was performed upon TB treatment initiation, after two months and at completion of treatment. To examine the relation between mycobacterial clearance in hosts and changes in T-cell immune-profiles, the results of these analysis were compared in negative and positive sputum culture conversion patients after two months of treatment.

## Results

### Study design and analysis strategy

Between May 2019 and July 2020, 144 cell samples collected from 22 adult TB patients were analyzed (Bangladesh, n=4 and Georgia, n=18; DS- and DR-TB, n=11 each) (Supp. Figure 1). Patient demographic, microbiological and clinical characteristics are available in Supp. Table 1. All patients achieved microbiological cure at the end of treatment, but were retrospectively classified into two response groups according to their *M. tuberculosis* culture status at T1 (after two months of treatment): fast converters (n=18; negative culture at T1 and T2) and slow converters (n=4; positive culture at T1 and negative culture at T2). Among the latter, three patients were treated for DS-TB and one for DR-TB.

An overview of the data collection and analysis process is shown in Figure 1. Briefly, data from all samples were clustered automatically into subsets of homogeneous phenotypes to provide a framework for analysis. Clusters were then color-coded and plotted onto a two-dimension map to create a visual reference used throughout the paper (Figure 2). On this basis, automatically detected clusters were first quantified and analyzed dynamically throughout treatment to identify median clusters abundance variations associated with treatment completion (Figure 3). Cluster phenotypes were deduced from marker expression heatmaps, and hierarchical clustering was applied based on marker expression (Figure 4). In a supervised manner, clusters of similar abundance changes and immunophenotypes were then re-grouped into larger subsets, in order to assess relevance of the detected abundance variations at the individual level, and consistency with manual gating (Figure 5). Finally, a cross sectional analysis was performed at T1 to identify which automatically detected clusters differentiated patients based on the microbiological response to the intensive phase of treatment (*Mtb* culture positivity at T1; Figures 6 and 7).

**Figure 1.**
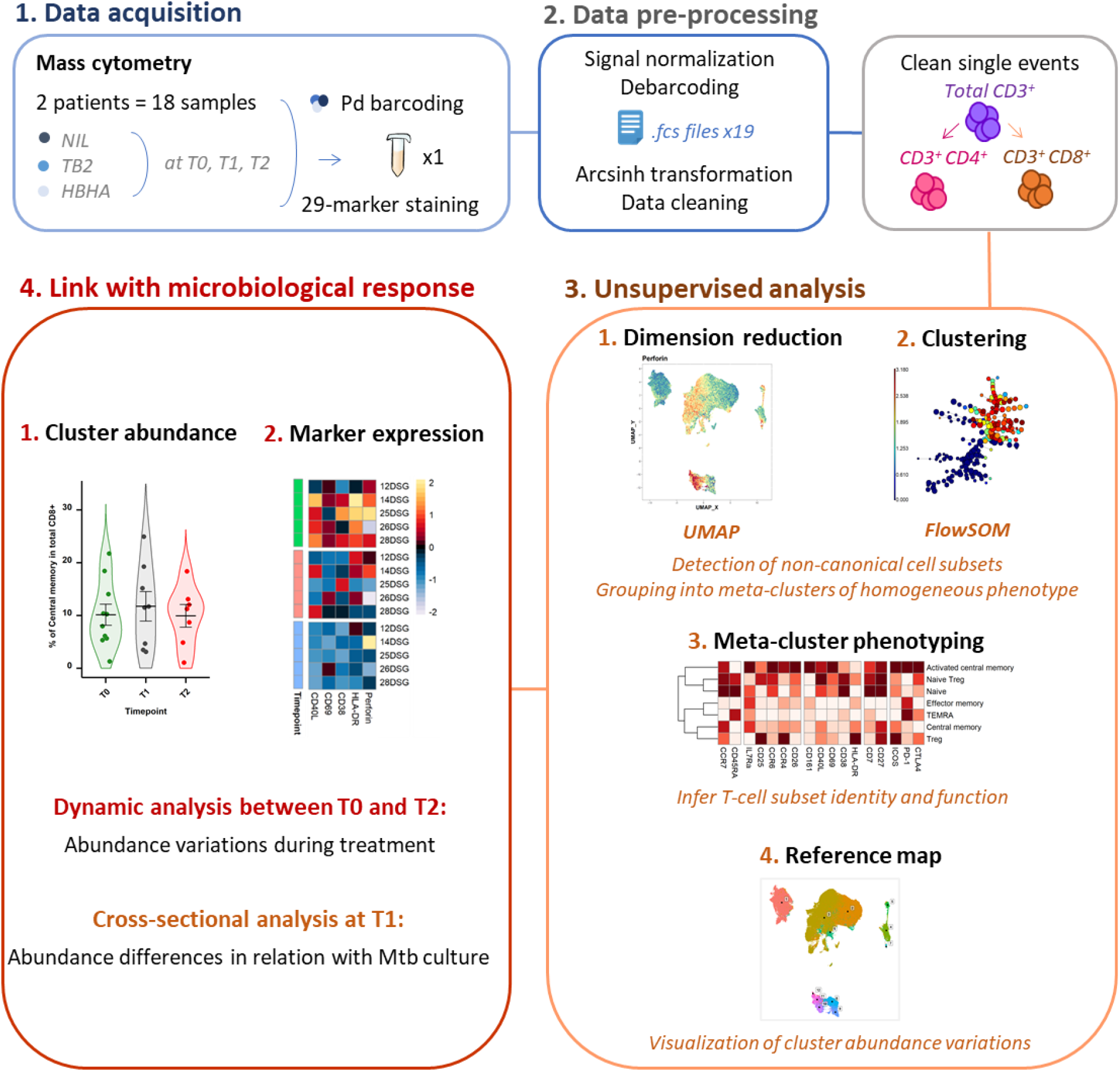
Experimental and analytical workflow. Peripheral whole blood samples were collected from active TB patients (n = 22) throughout treatment (T0: baseline. T1: T0 + 2 months. T2: end of treatment). After whole blood stimulation with *Mtb* antigens (TB2 and rmsHBHA) or with a negative control (NIL), total white blood cells were extracted. After palladium (Pd) barcoding for unique sample identification before multiplexing, T-cells were analyzed with a 29-marker mass cytometry panel. <u>Abbreviations:</u> TB2: Qiagen QuantiFERON TB2 tube (ESAT-6 + CFP-10 + undisclosed CD8^+^ T-cell stimulating peptide pool). rmsHBHA: recombinant heparin-binding hemagglutinin obtained in *Mycobacterium smegmatis.* UMAP: Uniform Manifold Approximation and Projection. FlowSOM: self-organizing map.

**Figure 2.**
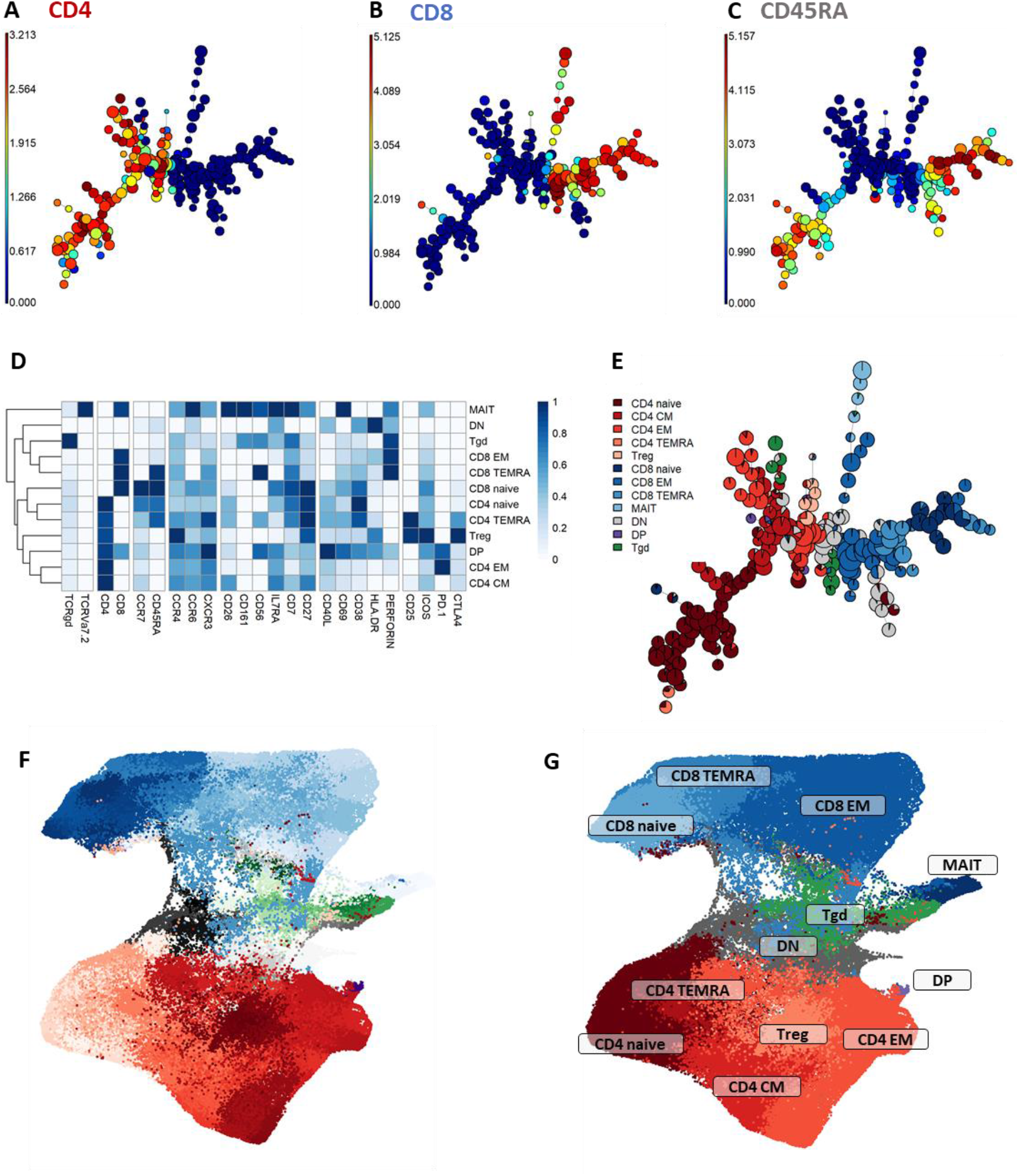
Peripheral CD3^+^ T-cell unsupervised clustering and phenotyping. **A to E. FlowSOM automated clustering.** The surface expression of lineage markers used for FlowSOM calculations was visualized in all CD3^+^ events (201,000 events from equally down-sampled files) regardless of timepoint or stimulation. FlowSOM enabled automated repartition of CD3^+^ events into 196 clusters according to the surface expression of selected lineage markers such as CD4 (**A**), CD8 (**B**), and CD45RA (**C**). Scales indicate arcsinh-transformed mass signal values. Clusters were automatically grouped into 18 meta-clusters of homogeneous phenotype, which were assembled into 12 canonical T-cell subpopulations in a supervised manner after meta-cluster phenotyping. This was performed with heatmap visualization of normalized, arcsinh-transformed median mass signal values for each surface marker (**D**). The proportions of the resulting T-cell subpopulations were visualized on the initial FlowSOM minimum spanning tree to control phenotyping consistency (**E**). **F and G. Reference mapping.** Dimension reduction was performed with UMAP and overlayed with automatically determined FlowSOM clusters (**F**) and meta-clusters (**G**) to generate a phenotype reference map. Cluster labels were not displayed for legibility. <u>Abbreviations:</u> CM: central memory. DN: double-negative CD4^-^CD8^-^. DN: double-positive CD4^+^ CD8^+^. EM: effector memory. MAIT: mucosal associated invariant T-cells. Tgd: gamma delta T-cells. Treg: T-regulators. TEMRA: terminally differentiated effectors re-expressing CD45RA.

**Figure 3.**
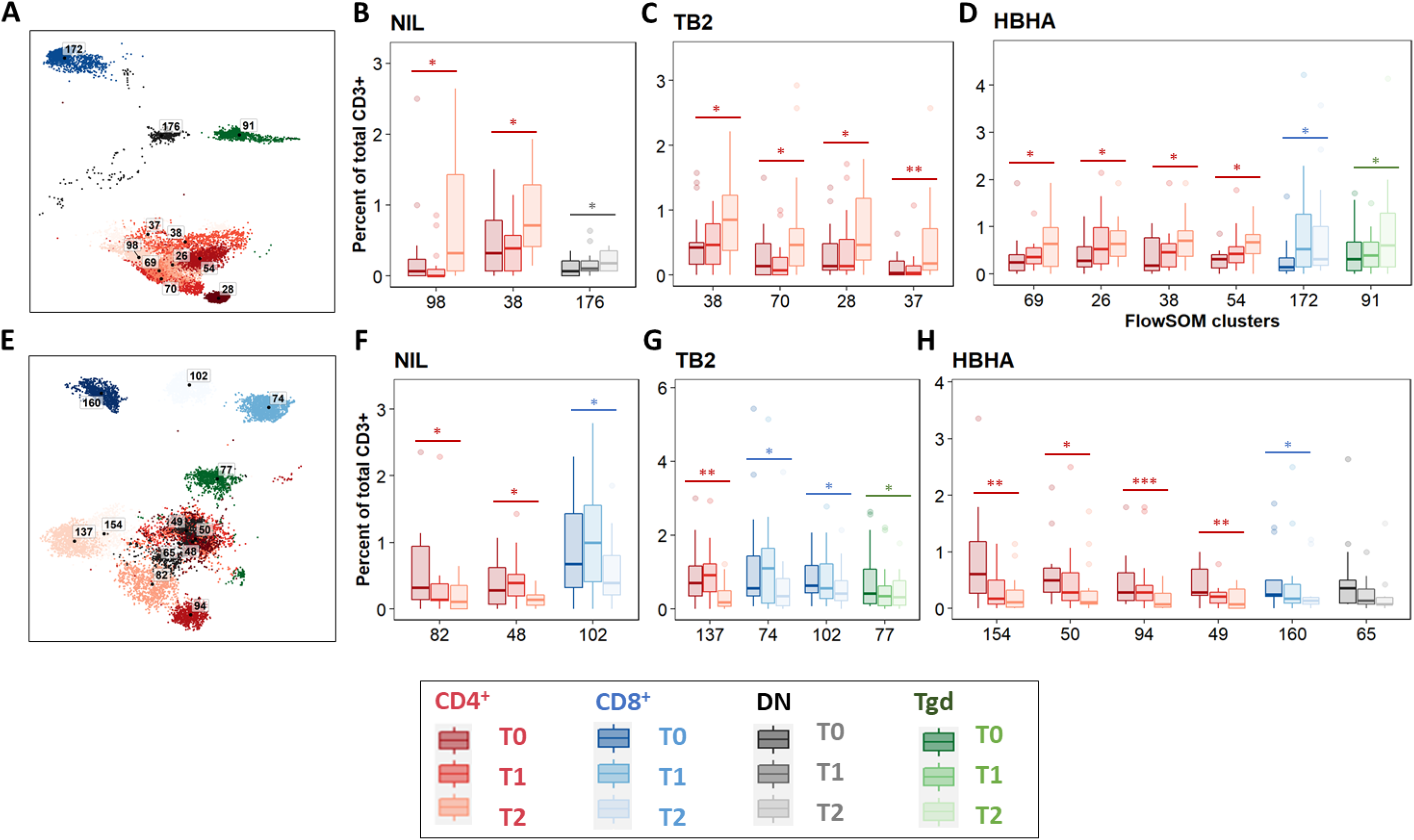
Significant abundance changes in non-canonical T-cell subsets throughout TB treatment. FlowSOM cluster abundance was analyzed over time in unstimulated or *Mtb*-stimulated samples (TB2 or rmsHBHA). Only clusters within which significant abundance changes were detected were displayed. Number of matched data points per timepoint for all panels: NIL: n = 16. TB2: n = 18. rmsHBHA: n = 14. Data are represented as medians + interquartile range. **A to D. Significantly increased clusters at treatment completion (T2) compared to treatment initiation (T0).** Clusters within which a significant increase was detected between T0 and T2 were visualized on the reference UMAP shown in Figure 3 (**A**). Cluster abundance quantification was was performed in unstimulated (**B**), TB2-stimulated (C) or rmsHBHA-stimulated samples (**D**). **E to H. Significantly decreased clusters at treatment completion (T2) compared to treatment initiation (T0).** Mapping (**E**) and abundance quantification of clusters which increased between T0 and T2 in unstimulated (**F**), TB2-stimulated (**G**) or rmsHBHA-stimulated samples (**H**). <u>Abbreviations:</u> DN: double negative CD4^-^ CD8^-^. Tgd: gamma delta T-cells. <u>Statistical analysis:</u> Friedman rank sum test and Wilcoxon-Nemenyi-Thompson post-hoc for pairwise comparisons between non-independent observations at T0, T1, and T2. *: p<0.05. **: p<0.01. ***: p<0.001. Exact p-values and test statistics are available in Supp. Table 3.

**Figure 4.**
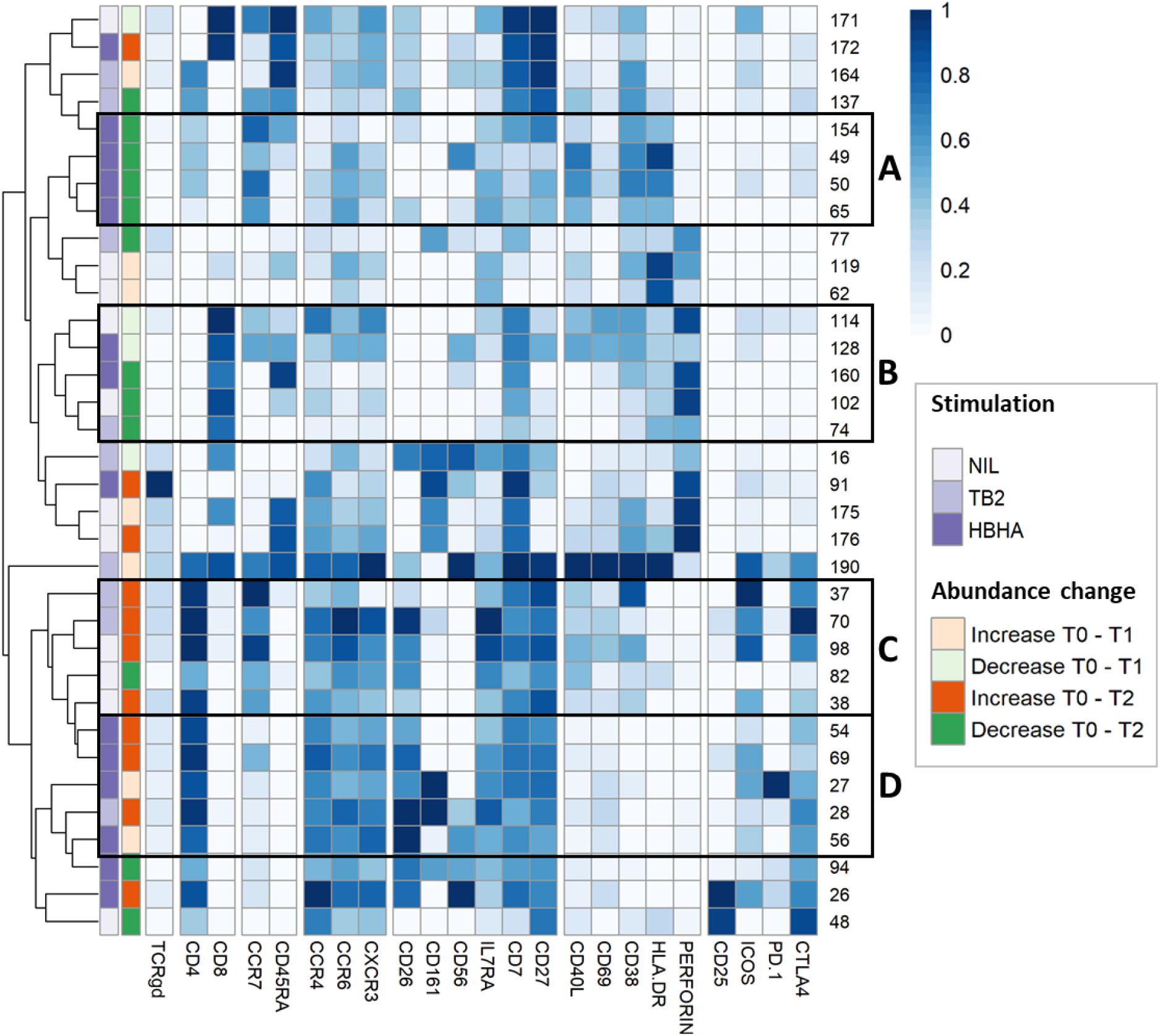
In-depth phenotyping shows differential involvement of effector and memory T-cells in antigen-driven cluster abundance changes during TB treatment. Mean marker expression levels were visualized using heatmapping for cell cluster which increased (orange color code) or decreased (green color code) throughout treatment. Each line represents one cell cluster. Scales indicate normalized mass signal intensity. Hierarchical clustering was performed based on marker expression levels, regrouping cell clusters of similar immunophenotypes. Black rectangles annotated from A to D indicate cell cluster subgroups with both similar immunophenotypes and abundance changes.

**Figure 5.**
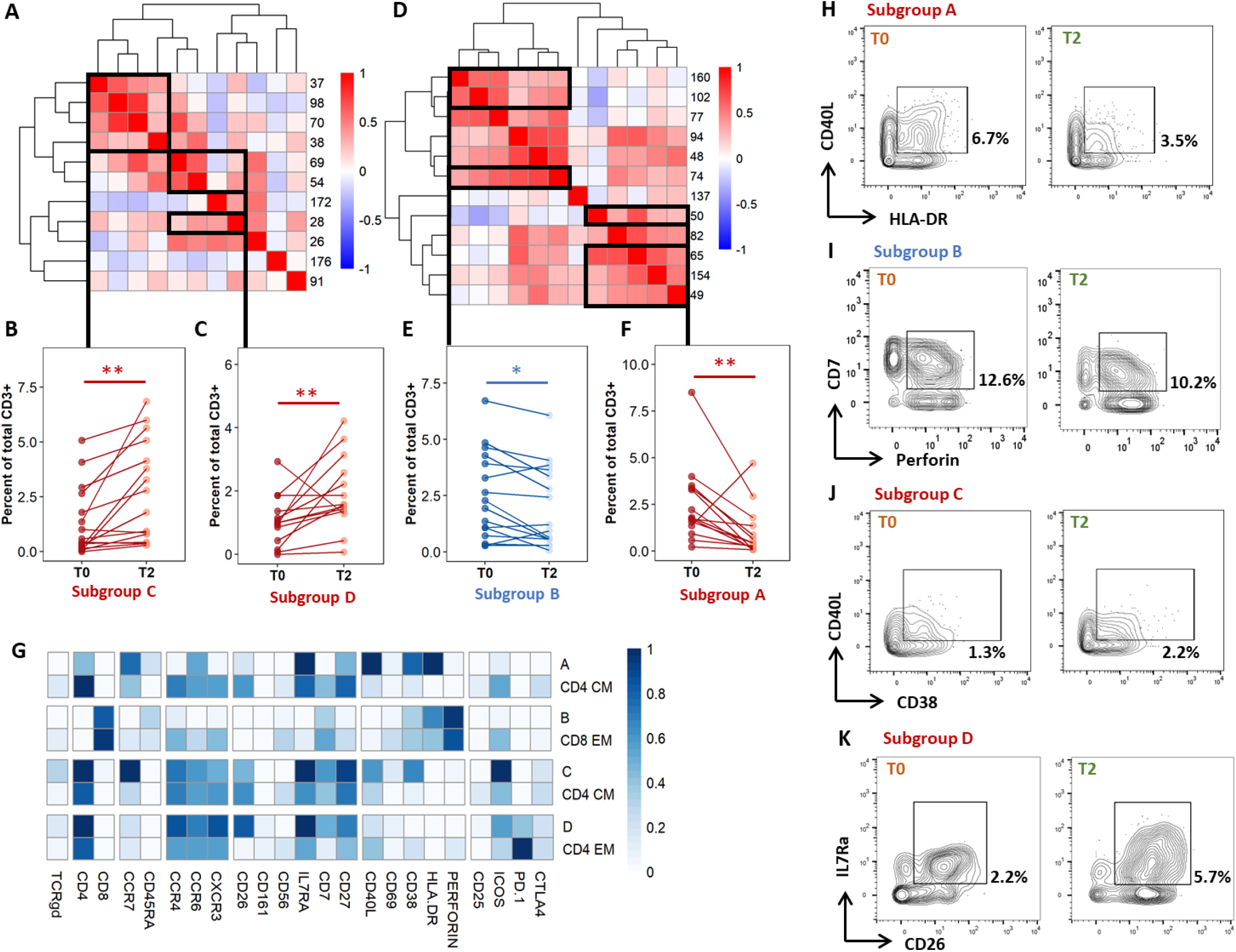
Individual immunoprofiling confirms differential abundance of correlated subsets in cured patients after treatment. Cluster were stratified by type of significant abundance change: enrichment (**A to C**) or depletion (**D to F**) after treatment completion. **A and D.** Pearson’s correlations were calculated on cluster abundance at T0 and displayed on a heatmap with hierarchical clustering. Clusters with similar immunophenotypes (Figures 3 and 4) and positive correlation coefficients were grouped. Estimates of effect sizes are in Supp. Tables 4 and 5. **B, C, E, F.** The abundance of each subgroup was visualized. Each dot represents data for one patient. <u>Statistical analysis:</u> Friedman rank sum test. *: p<0.05. **: p<0.01. Subgroup A: data from rmsHBHA samples (n =14), clusters 49, 50, 65, 154; p = 0.0013, Friedman’s Chi-Square (Fchisq) = 10.3. Subgroup B: data from unstimulated samples (n =16), clusters 74, 102, 160; p = 0.020, Fchisq = 5.4. Subgroup C: data from unstimulated samples, clusters 37, 38, 70, 98; p = 0.0027, Fchisq = 9. Subgroup D: data from rmsHBHA samples, clusters 28, 54, 69; p = 0.0023, Fchisq = 9.3. **F.** For each subgroup, normalized mean marker expression levels were compared with similar T-cell subsets. **G to K.** Manual gating analysis was performed to verify unsupervised results (representative plots, 500 to 1,000 events). Numbers indicate the percentage of gated cells among total CD3^+^ cells. Subgroup A: CD4^+^CCR7^+^CD45RA^-^CCR6^+^IL7Ra^+^CD27^+^CD40L^+^CD38^+^HLA-DR^+^. Subgroup B: CD8^+^CCR7^-^CD45RA^-^CD7^+^ Perforin^+^. Subgroup C: CD4^+^CCR7 CD45RA^-^CCR4^+^CCR6^+^CXCR3^+^CD26^+^IL7Ra^+^CD7^+^CD27^+^ CD40L^+^CD38^+^. Subgroup D: CD4^+^CCR7^-^CD45RA^-^CCR4^+^CCR6^+^ CXCR3^+^CD26^+^IL7Ra^+^CD7 CD27^+^CD40L^+^CD38^+^ HLA-DR^-^.

**Figure 6.**
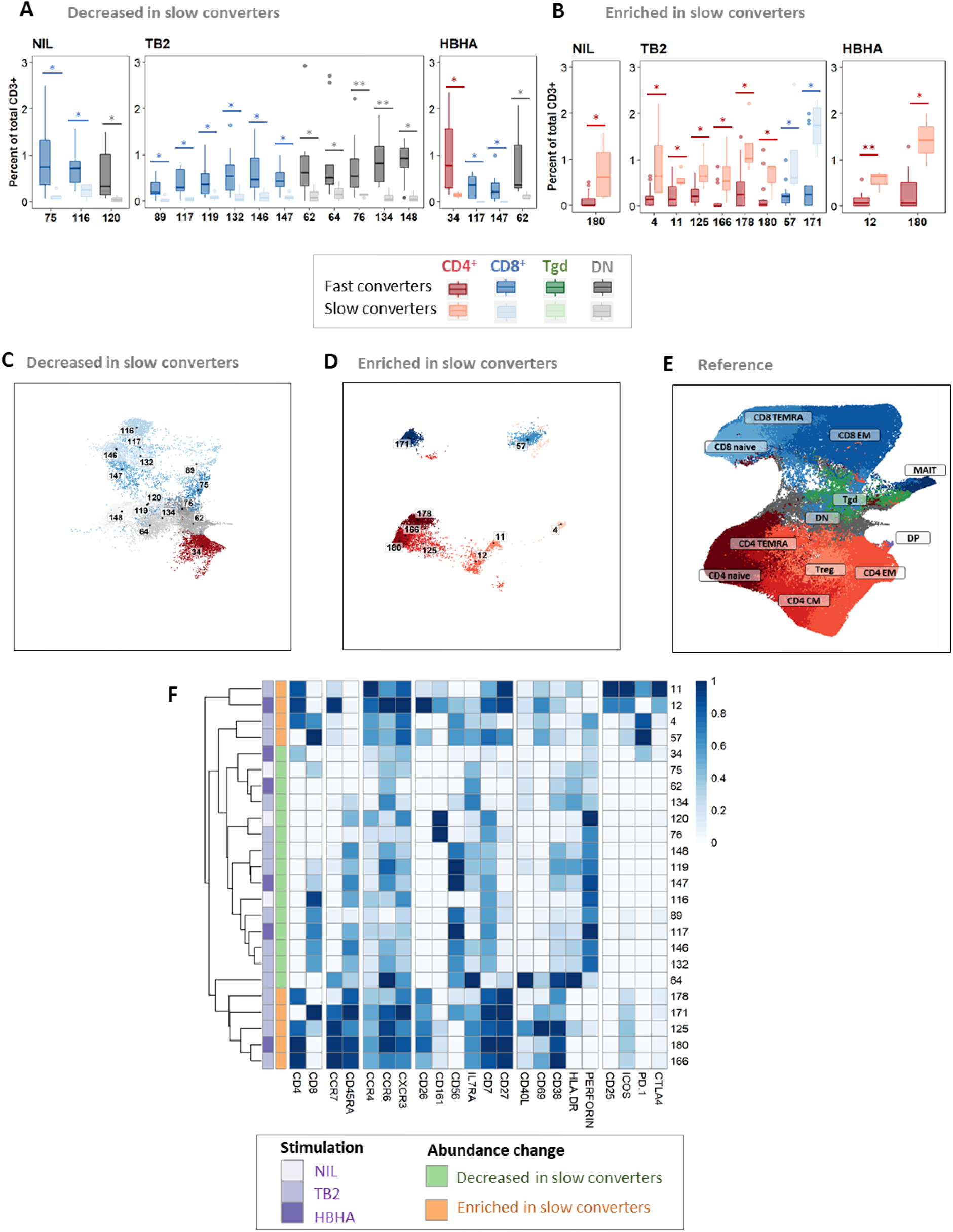
Patients with slow microbiological culture conversion show decreased cytotoxic CD8^+^ and γδ enriched CD4^+^ naïve T-cell subsets before treatment initiation and after two months of treatment compared to fast converters. Fast converters (n = 18) were defined as patients with permanently negative *M. tuberculosis* culture after two months of treatment (T1), whereas slow converters (n = 4) were defined as patients with persistently positive cultures at T1. The abundance of all FlowSOM clusters at baseline was compared between fast and slow converters. CD4^+^ clusters were represented in red, CD8^+^ clusters in blue, and γδ T-cell clusters in green. Clusters which were significantly decreased (**A** and **C**) or enriched (**B** and **D**) at T1 in slow converters compared to fast converters were compared to the reference UMAP (**E**). Normalized, arcsinh-transformed mean marker expression levels were visualized (**F**). Each row represents one cluster. Scales indicate normalized mass signal intensity. Boxplot data represent medians + interquartile range. <u>Statistical analysis:</u> Only clusters within which significant differences were detected were represented. Significance threshold: p<0.035 (Mann-Whitney U test). *: p<0.031. **: p<0.001. Exact p-values and test statistics are available in Supp. Table 6.

**Figure 7.**
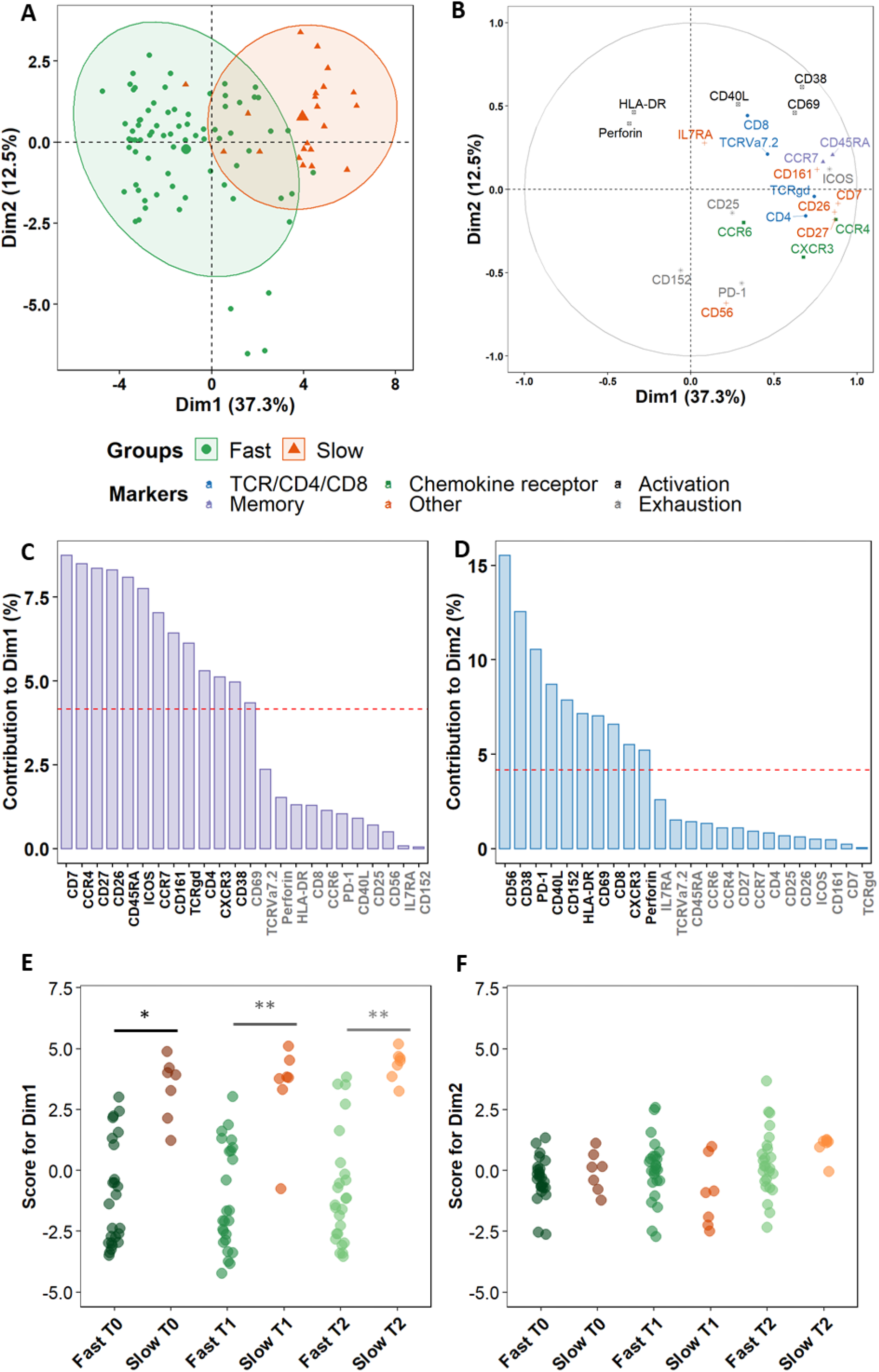
Non-lineage markers discriminate slow and fast responders within differentially abundant subsets. Principal Component Analysis (PCA) was performed on marker expression data from the clusters identified in Figure 6, within 96 *Mtb*-stimulated samples matched at T0, T1, and T2 (TB2: 54 samples; rmsHBHA: 42 samples; see Supp. Table 1 for sample number details). **A.** Explanation of the variance between fast converters (25 samples at each timepoint) and slow converters (7 samples at each timepoint). Axes represent the principal components 1 (Dimension 1, Dim1) and 2 (Dim2). Percentages indicate their contribution to the total observed variance. Axis values represent individual PCA scores. Concentration ellipses correspond to 90% data coverage. **B.** Contribution of cellular markers to the variance described by Dim1 and Dim2. Axis values represent marker PCA scores. Color codes represent broad marker functions. **C and D.** Quantification of panel B for Dim1 (**C**) and Dim2 (**D**). Contributions of each marker are expressed as a percentage of the dimensions. The dashed line corresponds to the expected reference value if each marker contributed uniformly to the variance. Markers indicated in gray are below this reference value. **E and F.** Distribution of individual PCA score values according to the culture conversion group at each timepoint, for Dim1 (**E**) and Dim2 (**F**). Wilcoxon Rank Sum Test. ***: p<0.001. **: p<0.001. Exact p-values and test statistics are in Supp. Table 7.

### Overall analysis of peripheral T lymphocyte subset abundance changes throughout TB treatment

First, a phenotype analysis was performed to identify the main expected T-cell subpopulations. As no apparent difference was seen in UMAP structures within samples from the different timepoints and stimulation conditions despite some marker expression differences between stimulation conditions (Supp. Figure 2; exact p-values and test statistics in Supp. Table 2), we performed the phenotype analysis on all single CD3^+^ events. The purpose of this study was not to compare the stimulations, but rather to use them to uncover clusters that might be associated with treatment response and that would not be visible in unstimulated samples. FlowSOM automated clustering was performed on CD3^+^ events, revealing a total of 196 automatically detected clusters (Figure 2.A to 2.C). They were automatically grouped into 18 meta-clusters, which were assembled into 12 canonical T-cell subpopulations in a supervised manner (Figure 2.D and 2.E). FlowSOM clusters and meta-clusters were then visualized on the initial UMAP to create a reference map of all automatically detected T-cell subsets (Figure 2.F and 2.G). To initiate the abundance analysis, variations of the main T-cell subpopulations throughout treatment were then studied using a stratification according to each stimulation condition. No significant change in the proportion of total CD4^+^, CD8^+^, γδ, double negative (DN, CD4^-^ CD8^-^) or double positive (DP, CD4^+^ CD8^+^) T-cells was observed throughout treatment in any stimulation condition (Supp. Figure 3). For all main studied subpopulations, no significant difference was observed between DS- and DR-TB patients (data not shown).

### Differential abundance of non-canonical T-cell subsets throughout TB treatment

To identify non-canonical T-cell subsets whose abundance changed throughout treatment, we calculated the percentage of each automatically determined FlowSOM cluster at each timepoint and in each stimulation condition. These clusters were then categorized into two groups: enriched or decreased after treatment completion. Abundance changes were studied between T0 and T1 and T0 and T2 to characterize the main clusters associated with response to treatment intensive phase and with treatment completion respectively. As these clusters represent non-canonical cell subpopulations, their frequencies among total CD3^+^ events were low (< 5% in most samples). Hence, the differences analyzed thereafter describe rare populations and warrant cautious analysis.

When comparing the reference UMAP (Figure 2.G) to the UMAP of clusters which were increased between T0 and T1 (Supp. Figure 4.A), we observed that they were either DN T-cells, or effector memory (EM) or terminally differentiated effectors re-expressing CD45RA (TEMRA) cells from both CD4^+^ and CD8^+^ subpopulations. In unstimulated samples, significant increases were detected within three clusters corresponding to CD8^+^ and DN T-cell subsets (Supp. Figure 4.B), whereas increases were detected in one CD4^+^ and one CD8^+^ cluster in TB2-stimulated samples (Supp. Figure 4.C) and only in CD4^+^ clusters in rmsHBHA samples (Supp. Figure 4.D). Clusters that decreased between T0 and T1 (Supp. Figure 4.E) were detected only within CD8^+^ EM and TEMRA cells in all stimulation conditions (Supp. Figure 4.F to 4.H). Between T0 and T2, 11 increased clusters were detected (Figure 3.A). They corresponded mostly (8/11 clusters, 73%) to CD4^+^ EM and CM subpopulations rather than naïve subsets, regardless of the stimulation condition (Figure 3.B. to 3.D.). One DN cluster was increased in unstimulated samples (Figure 3.B. as well as one CD8^+^ TEMRA cluster and one γδ T-cell cluster in rmsHBHA stimulated samples (Figure 3.D.). One CD4^+^ CM cluster (number 38) increased significantly in samples from all three stimulation conditions. Clusters which decreased between T0 and T2 were detected in one CD8^+^ EM and two CD8^+^ TEMRA subsets, and in seven clusters within CD4^+^ subpopulations in all three stimulation conditions (Figure 3.E to 3.H). Regarding the latter clusters, no clear trend was observed regarding memory subset compartmentalization, which suggests that the abundance decrease spared memory functions and rather affected CD4^+^ T-cells in general. One γδ and one DN T-cell cluster also decreased significantly within *Mtb-*stimulated samples (Figure 3.G. and 3.H.).

### Antigen-driven cluster abundance changes during TB treatment show involvement of effector and memory T-cells

To further refine patterns in functional marker expressions within increased or decreased clusters, we then performed a detailed phenotype analysis using marker expression heatmaps and hierarchical clustering (Figure 4). Four subgroups of cellular subsets of similar abundance changes and similar immunophenotypes were identified (labeled from A to D). Subgroup A included four CD4^+^ T-cell clusters with naive (n=2) and CM (n=2) phenotypes, which decreased from T0 to T2 in rmsHBHA-stimulated samples. Subgroup B included five CD8^+^ T-cell clusters that decreased throughout treatment, two of them between T0 and T1 and three of them between T0 and T2. Consistently with the above results (Figure 3.E.), the latter were either EM or TEMRA cells, with low CD45RA levels and intermediate levels of perforin. The other two clusters were naïve clusters with low CCR7, CD45RA, and CD27 expression levels.

In contrast, subgroup C and D included only CD4^+^ T-cell clusters, most of which (70%, 7/10) increased between T0 and T2. Subgroup C consisted in five clusters exhibiting a CM phenotype and expressing activation markers, detected in unstimulated and TB2-stimulated samples. Subgroup D clusters were detected in *Mtb*-stimulated samples (3 in rmsHBHA and 2 in TB2) and had an EM phenotype, except for cluster 69 that had a CM phenotype with low levels of CCR7. These clusters co-expressed CD26, IL7Ra, CD7 and CD27. They were characterized by an absence of activation marker expression and an enhanced expression of exhaustion markers, in particular CTLA-4 and PD-1. Overall, we observed antigen-driven T-cell subset abundance changes between T0 and T2. In TB2 and rmsHBHA samples, CD4^+^ EM clusters mostly increased, while CD8^+^ EM clusters mostly decreased.

### Individual profiling confirms abundance changes in phenotypically homogeneous, correlated subsets after treatment in cured patients

As the differentially abundant clusters identified above accounted for a small fraction of CD3^+^ T-cells (<1%), we intended to identify the largest possible subsets of phenotypically homogeneous cells within which a significant abundance change was detectable (Figure 5). Within the subgroups of similar immunophenotypes and abundance change identified in Figures 3 and 4, we performed correlation analyses at baseline and pooled the best correlated clusters together within the subgroups identified in Figure 4 (Figure 5.A and 5.D). We then visualized the individual abundance change of these pooled subsets before and after treatment completion in cured patients (Figure 5.B-C and 5.D-E). Within rmsHBHA samples, a decrease in subgroup A and an increase in subgroup D were both detected in 93% (13/14) of cured participants (Table 1). Within unstimulated samples, a decrease in subgroup B and an increase in subgroup C were recorded in 81% (13/16) and 88% (14/16) of patients respectively. This confirmed that the median trends observed previously were maintained individually in most patients. Finally, we visualized the immunophenotypes of these four subgroups of interest in comparison to cells from similar subpopulations which were not associated to cure (Figure 5.F). Subgroup A and subgroup C corresponded to CD4^+^ CM cells expressing CCR6, IL7Ra, CD27, and activation markers (CD40L, CD38). However, cells within subgroup A expressed HLA-DR while subgroup C did not; in addition, cells from subgroup C expressed high levels of CD26, as well as CCR4, CXCR3, and CD7. Subgroup B corresponded to CD8^+^ CD7^+^ Perforin^+^ EM cells. Subgroup D corresponded CD4^+^ EM cells expressing high levels of CD26, as well as CCR4, CCR6, CXCR3, IL7Ra, CD7, and CD27. We then confirmed these findings by manually gating the identified subpopulations and comparing the percentages at T0 and T2 (Figure 5.G-K, representative dot plots).

**Table 1.** Selected subset abundance changes before and after treatment completion.

| Sample | Abundance between T0 and T2 (% , N) |
| --- | --- |
| **Subset A decreased** |  |
| NIL (n=16) | 62% (10) |
| TB2 (n=18) | 67% (12) |
| rmsHBHA (n=14) | **93% (13)** |
| **Subset B decreased** |  |
| NIL | **81% (13)** |
| TB2 | 72% (13) |
| rmsHBHA | 71% (10) |
| **Subset C increased** |  |
| NIL | **88% (14)** |
| TB2 | 72% (13) |
| rmsHBHA | 57% (8) |
| **Subset D increased** |  |
| NIL | 69% (11) |
| TB2 | 78% (14) |
| rmsHBHA | **93% (13)** |

Footnotes: these data were obtained from Figure 5.

### Patients with persistent positive cultures at T1 show decreased peripheral CD8^+^ cytotoxic subsets and enriched peripheral CD4^+^ naïve subsets throughout treatment compared to patients with negative cultures at T1

Then, we aimed to detect a cellular signature associated with mycobacterial conversion. To do so, we analyzed individual cluster abundance in slow *vs*. fast converters throughout treatment. At T0, T1, and T2, respectively 21, 24, and 21 clusters with significantly different abundance in slow converters compared to fast converters were detected (quantification in Supp. Fig. 5). After phenotyping, the proportions of the main T-cell subpopulation phenotypes in each group of enriched or decreased clusters at T0, T1, and T2 were calculated and summarized in Table 2. Before treatment initiation, of 21 clusters with different abundance, 18 (86%) were decreased (Supp. Figure 5.A) and three (14%) were enriched (Supp. Figure 5.B) in slow compared to fast converters. Clusters which were under-represented in slow converters corresponded mostly to DN, γδ, and CD8^+^ T-cells (77%, 13/18 clusters), specifically γδ and CD8^+^ EM T-cell subpopulations (38%, 5/13 each); in addition, a majority of these clusters was perforin^+^ (67%, 12/18) (Supp. Figure 6.A). In contrast, the three enriched clusters were naive CD4^+^ and CD8^+^ T-cells, as well as one CD8^+^ TEMRA subset.

**Table 2.** Proportions of the main T-cell subpopulations within enriched or decreased subsets in slow converters compared to fast converters.

|  | T0 (21 clusters) |  | T1 (24 clusters) |  | T2 (21 clusters) |  |
| --- | --- | --- | --- | --- | --- | --- |
| **Abundance in slow vs. fast converters** | **Decreased**<br>**86% (18)** | **Enriched**<br>**14% (3)** | **Decreased**<br>**62% (15)** | **Enriched**<br>**38% (9)** | **Decreased**<br>**52% (11)** | **Enriched**<br>**48% (10)** |
| **Total CD8<sup>+</sup> and $\gamma\delta$** | **72% (13)** | **67% (2)** | **53% (8)** | **22% (2)** | **36% (4)** | **20% (2)** |
| $\gamma\delta$ T-cells | 38 (5) | - | - | - | - | - |
| CD8<sup>+</sup> TEMRA | 24 (3) | 50 (1) | 75 (6) | - | - | - |
| CD8<sup>+</sup> EM | 38 (5) | - | 25 (2) | 50 (1) | 100 (4) | - |
| CD8<sup>+</sup> naïve | - | 50 (1) | - | 50 (1) | - | 100 (2) |
| **Total CD4<sup>+</sup>** | **11% (2)** | **33% (1)** | **7% (1)** | **78% (7)** | **36% (4)** | **80% (8)** |
| CD4<sup>+</sup> TEMRA | - | - | - | 14 (1) | - | - |
| CD4<sup>+</sup> EM | 50 (1) | - | 100 (1) | 29 (2) | 100 (4) | - |
| CD4<sup>+</sup> CM | 50 (1) | - | - | 14 (1) | - | 38 (3) |
| CD4<sup>+</sup> naïve | - | 100 (1) | - | 43 (3) | - | 62 (5) |
| **Total DN** | **17% (3)** | **0** | **40% (6)** | **0** | **27% (3)** | **0** |

At T1, of 24 clusters with significantly different abundance between slow and fast converters, 15 (62%) were decreased (Figure 6.A and 6.C) and 9 (38%) were enriched in slow converters (Figure 6.B and 6.D). These clusters were mostly detected in TB2-stimulated samples (63%; 15/24 clusters). Comparison to the reference UMAP (Figure 6.E) and hierarchical clustering (Figure 6.F) indicated that enriched and decreased subsets respectively had similar immunophenotypes. Clusters which were under-represented at T1 in slow converters were mostly perforin^+^ cells (67%, 10/15 clusters); mostly CD8^+^ TEMRA and DN T-cell phenotypes were represented (40%, 6/15 clusters respectively). In contrast, enriched clusters comprised a majority of CD4^+^ T-cells (78%, 7/9 clusters), with predominantly naïve phenotypes (45%, 3/7). One CD8^+^ naive and one CD8^+^ EM cluster were also enriched in slow converters at T1, with the latter expressing ICOS.

After treatment completion, of 21 clusters with significantly different abundance between slow and fast converters, 11 (52%) were decreased (Supp. Figure 5.C) and 10 (48%) were enriched in slow converters (Supp. Figure 5.D). The immunophenotype profile at T2 was similar to that of T1 for the enriched subsets: a majority of ICOS^+^ CD4^+^ naïve T-cell subsets (50%, 5/10) were detected, as well as two CD8^+^ naïve clusters (Supp. Figure 6.B). Regarding the decreased subsets, no specific phenotype polarization was observed, and clusters were detected within diverse subsets (four CD8^+^ EM clusters, four CD4^+^ EM clusters, and three DN T-cells clusters). Similarly to the T1 immune profile, all of the above clusters were mostly detected in TB2-stimulated samples (67%, 14/21 clusters).

### Maturation markers and chemokine receptors, rather than activation or cytotoxic markers, discriminate slow from fast converters during treatment

Finally, we sought to assess more precisely which combinations of cellular markers were the most involved in the discrimination between fast and slow converters within the clusters identified in the prior section. A principal component analysis (PCA) was performed on marker expression data within these clusters. As a higher number of differentially abundant clusters had been detected in *Mtb*-stimulated samples than in unstimulated samples during treatment (T1 and T2), and because a complete overlap between the PCA profiles of fast and slow converters was observed in unstimulated samples, we focused on *Mtb*-stimulated samples (TB2 and rmsHBHA). PCA profiles were mostly separated when split by culture conversion group (Figure 7.a). Dimension 1 (Dim1) explained 37.3% of the total observed variance, versus 12.5% for Dim2. The main markers accounting for variance described by Dim1 were markers of memory subset definition (CCR7 and CD45RA), lineage (CD4 and TCRγδ), maturation (CD27 and CD7), chemokine receptors (CCR4 and to a lesser extent CXCR3) or other receptors or costimulatory molecules (*e.g.*, CD26, CD161) (Figure 7.B. and 7.C). In contrast, variance described by Dim2 was mostly explained by cytotoxicity (Perforin, CD56, CD8), activation (CD38, CD40L, CD69), or exhaustion markers (CD152, PD-1) (Figure 7.B and 7.D). The PCA scores were significantly higher in slow converters than in fast converters at all timepoints for Dim1 (Figure 7.E), indicating that the immune profile of slow converters was more correlated to Dim1 than that of fast converters regardless of the timepoint. In contrast, no significant differences were detected at the end of treatment (T2) for Dim2 (Figure 7.F). When comparing these results with PCA analyses performed on total CD3^+^ T-cells, fast and slow converter profiles were less separated, but similar marker involvement was observed in Dim1 and Dim2 respectively (Supp. Figure 7).

## Discussion

In a population of adults treated for TB, we observed a shift towards more differentiated profiles among peripheral CD8^+^ and CD4^+^ T-cell subsets driven by the timing of *Mtb* culture conversion, using a high-dimensional single cell approach after stimulation with standardized, IVD-level TB2 antigens. In particular, differentiated CD8^+^ cytotoxic effector subsets were under-represented in positive-versus negative-sputum culture patients after two months of treatment.

Over the course of TB treatment, we observed as a general trend that non-canonical subsets within CM CD4^+^ and TEMRA CD8^+^ populations increased, whereas naïve CD4^+^ and naïve/EM CD8^+^ subsets decreased. This is consistent with prior works addressing T-cell differentiation and T-cell memory subsets during TB treatment (Marriott et al., 2018; Chiacchio et al., 2014; Wang et al., 2010). *Mtb*-specific CD4^+^ EM T-cells have been associated with active TB disease, whereas CM T-cells have been associated to latency and increased upon treatment (Petruccioli et al., 2013; Goletti et al., 2006a). In *Mtb*-specific CD8^+^ T-cells, an overall decrease in peripheral blood (Nyendak et al., 2013) and a decrease in CM cells (Axelsson-Robertson et al., 2015b) have been documented after treatment. In contrast, the central result of this study was to distinguish negative-from positive-sputum culture patients at two months, whether infected with a DS-or DR-*Mtb* strain, through differential peripheral T-cell populations. When retrospectively analyzing the T-cell profiles of fast and slow converters at diagnosis, a pre-existing difference in percentages of cytotoxic EM CD8^+^ T-cell subpopulations was already observed. After two months of treatment, this trend shifted into an under-representation of CD8^+^ TEMRA, which persisted after cure. These changes were revealed upon stimulation with QFT-P TB2 antigenic peptide pools. Although many studies characterizing T-cell subsets during treatment have clearly underlined the importance of *Mtb*-specific CD4^+^ T-cells (Riou et al., 2014; Ahmed et al., 2018b; Riou et al., 2020), less is known about the role of CD8^+^ T-cells in TB resolution and the most appropriate epitopes to study them in this context (Lewinsohn et al., 2017; Chiacchio et al., 2018). Yet, effector CD8^+^ T-cells are known to secrete cytolytic and antimicrobial factors that kill *Mtb*-infected macrophages *in vitro* (Serbina et al., 2000), inhibit *Mtb* growth (Lewinsohn et al., 2017), and are required for long-term infection control in mice (Lin and Flynn, 2015) and humans (Bruns et al., 2009); perforin production by CD8^+^ T-cells is also higher in treated than in untreated TB patients (Jiang et al., 2017). In addition, a 2012 study by Rozot and colleagues had associated *Mtb*-specific TEMRA CD8^+^ T cells to LTBI and EM cells to active TB (Rozot et al., 2013). Here, although we cannot establish causality, a lower peripheral CD8^+^ TEMRA subset abundance may be associated with slower mycobacterial culture conversion. In relation with abundance changes during treatment, our study hints that the CD8^+^ T-cell phenotype shift occurring during TB treatment would be delayed in patients with slower microbiological conversion. Consistently, it has been shown that CD8^+^ response importantly contributed to the control of other granulomatous infections such as *Brucella* (Durward et al., 2012). Regarding CD4^+^ T-cells, naïve subsets were over-represented in slow converters, which suggests a delayed differentiation within the CD4^+^ compartment as well. Previous work has shown that the IFN-γ/IL-2/TNF-α functional profile of *Mtb*-specific CD4^+^ T-cells, which is key in anti-TB immunity (Chiacchio et al., 2017), was correlated with their degree of differentiation (Riou et al., 2017). Taken together, these results support the hypothesis that CD4^+^ and CD8^+^ T-cell responses should be monitored together during TB treatment, as successful mycobacterial clearance involves CD8^+^ T-cell effectors, which in turn require CD4^+^ T-cell involvement (Grotzke and Lewinsohn, 2005).

Although the aim of this study was not to compare stimulation conditions, but to use them to uncover cell clusters, our results suggest that the abundance changes observed throughout treatment are antigen-driven. This adds to previous work highlighting differential *Mtb*-specific CD8^+^ T-cells marker profiles according to the nature of the antigen stimulation (Axelsson-Robertson et al., 2015a). We used QFT-P TB2, which elicits cytotoxic CD8^+^ responses in addition to ESAT-6/CFP-10-induced CD4^+^ responses (Petruccioli et al., 2016), as well as rmsHBHA, a recombinant *Mtb* protein exposing many different epitopes. The latter was included because the IFN-γ response to HBHA, to which both CD4^+^ and CD8^+^ cells participate (Masungi et al., 2002), is impaired in active TB patients and restored during treatment (Chedid et al., 2021; Sali et al., 2018; De Maio et al., 2018). Here, changes during treatment in CD8^+^, CD4^+^, DN, and γδ T-cell subsets were detectable within unstimulated and TB2 samples, consistently with previous works (Petruccioli et al., 2016). In contrast, in rmsHBHA-stimulated samples, significant abundance changes were mostly detected within CD4^+^ T-cells, suggesting a preferential CD4^+^ T-cell response to HBHA epitopes during treatment. This indicates that antigen-driven changes during the response to *Mtb* are part of a complex process involving a variety of different epitopes (Axelsson-Robertson et al., 2015b) that induce responses from phenotypically diverse T-cell subsets (Axelsson-Robertson et al., 2015a), despite well-described immunodominance features. Our results confirm that a major stake in discovering blood-based immune signatures of mycobacterial sterilization lies in finding the appropriate epitopes.

Finally, our study enabled profiling of non-lineage markers. A CXCR3^+^ CCR6^+^ CD27^+^ CD4^+^ EM subset was increased in cured patients compared to pre-treatment, corresponding to a subset enriched in Th1/Th17 cells (Kim et al., 2001; Acosta-Rodriguez et al., 2007). Consistently with previous work on LTBI (Lindestam Arlehamn et al., 2013), this suggests that an increase in these cells upon cure might be associated with infection control. Compared to the other CD4^+^ EM cells, this subset displayed higher CD26 and IL7Ra expression. CD26 participates in T-cell activation and proliferation (Klemann et al., 2016), and correlates with Th1-like responses (Ohnuma et al., 2008). In parallel, a significant decrease was also observed in a highly activated CCR6^+^ IL7Ra^+^ CD4^+^ CM subset, which expressed higher levels of CD40L, CD38, and HLA-DR than other CD4^+^ CM cells. Interestingly, an increase in another CD4^+^ CM subset – which differed from the latter because it expressed CD26 and CD27, but not HLA-DR – was observed simultaneously. This adds to previous works highlighting a decrease in CD38^+^ and HLA-DR^+^ *Mtb*-specific CD4^+^ T-cells in successfully treated TB patients (Ahmed et al., 2018a; Riou et al., 2020; Adekambi et al., 2015b). This suggests that upon TB treatment, differentiated Th1/Th17-like CD4^+^ subsets expressing high levels of CD26 and IL7Ra are enriched in peripheral blood, likely at the expense of less differentiated subsets expressing high levels of CD27 and CD38. Finally, principal components analysis showed that within the subpopulations that differentiated slow from fast converters during treatment, differentiation markers and chemokine receptors contributed to most of the variance, followed by activation and cytotoxicity markers. CD27, CD26, and CCR4 were among the markers which best discriminated fast and slow responders, consistently with prior studies associating CD27 and CCR4 expression in *Mtb*-specific CD4^+^ T-cells with active TB compared to latent infection (Latorre et al., 2019). HLA-DR and CD38 also contributed to a lesser extent, which adds to a recent study in which co-expression of CD27, HLA-DR, and CD38 on PPD-stimulated CD4^+^ T-cells stratified fast and slow responders without restriction to IFN-γ-producing cells (Vickers et al., 2020).

This descriptive study has limitations. The number of patients included was low, resulting in few slow converters, consistently with treated TB course (15 to 20% of slow culture converters). In addition, the presence of within-host *Mtb* isolate micro-diversity has been recently proven in patients treated for DS-TB without culture conversion after two months of well-conducted TB treatment (Genestet et al., 2021), suggesting that it could modulate the host response. We are currently conducting a larger validation study including DS-TB patients only, from whom *Mtb* isolates collected upon treatment initiation and at two months will be screened by whole genome sequencing. In addition, the analyses were not conducted on live cells, but on fixed, cryopreserved peripheral blood cells due to the design of the study using samples collected in lower-income, high TB prevalence settings. For the same reason, the study was conducted on peripheral blood, while the main infectious focus of TB is in the lungs. In addition, since the study required to IGRAs to be performed on the same blood samples prior to cell cryopreservation (Chedid et al., 2021), we did not perform intracellular cytokine staining. Hence, the integrality of the observed cell phenotype changes may not be associated with *Mtb*-specific responses. However, whether the bulk of anti-TB response relies purely on *Mtb*-specific cells is debated. Given the complexity of the immune response to TB, cellular and molecular interactions are likely to occur between *Mtb*-specific and non-specific subpopulations during mycobacterial clearance, and hence influence the overall T-cell profiles. In addition, the hypothesis that T-cells specific for immunodominant epitopes actually recognize *Mtb*-infected cells has been challenged by studies on mouse models (Patankar et al., 2020), protective immunity post-BCG vaccination(Kagina et al., 2010), and failures of vaccine candidates based on immunodominant antigens (Moguche et al., 2017).

These limitations are linked to the “bench to bedside” approach adopted in our study. They reflect the reality of the needs for novel TB management tools: accessible samples, simple experimental process, straightforward output. Here, we captured the complexity of T-cell profiles during treatment and narrowed it down to subpopulations of interest associated with cure at the individual level. Although mass cytometry requires complex equipment, experiments, and analyses, we have shown that relevant T-cell profiles could be identified in cryopreserved samples, obtained from small blood volumes, using manual gating analyses and a smaller number of core markers. Future validation studies might confirm the relevancy of simpler phenotypic signatures translatable in primary care settings. Importantly, our study revealed T-cell populations discriminating patient status based on culture conversion, which has a dual impact: on TB management, to better characterize the phenotypes of T-cells involved in TB clearance; and on biomarker research, further supporting that a diversity of epitopes is needed to fully disclose the spectrum of these cells. This work may help identify simpler prognostic biomarkers associated with mycobacterial clearance and the antigens appropriate for their discovery.

## Materials and methods

### Experimental design

#### Study design and research objectives

This prospective cohort study was nested in a multicentered study coordinated by the Mérieux Foundation GABRIEL network (Chedid et al., 2020). The primary objective was to investigate the association between sputum culture sterilization during TB treatment and T-cell profiles obtained by high-dimensional phenotyping. The sample size was maximized based on availability of clinical samples. No prospective sample size calculations were performed.

#### Recruitment centers and ethical considerations

Recruitment centers were the National Center for Tuberculosis and Lung Disease (NTCLD) in Tbilisi, Georgia (approval of the Institutional Review Board of the NTCLD; IORG0009467); and the International Centre for Diarrhoeal Disease Research, Bangladesh (icddr,b) in Dhaka, Bangladesh (approval of the Research Review Committee and the Ethical Review Committee of icddr,b; PR-17076; Version No. 1.3; Version date: 04-01-2018). All participants provided written informed consent.

#### Cohort recruitment, patient follow-up, and clinical data collection

Patients were recruited if diagnosed with sputum culture confirmed pulmonary TB and older than 15 years old. Patients with HIV, immune deficiency, diabetes mellitus, and lost-to-follow-up were excluded. Detailed procedures for microbiological diagnosis, drug susceptibility testing, and treatment regimens are described elsewhere (Chedid et al., 2020). As antimicrobial resistance is a major challenge for TB management and treatment, both drug-susceptible (DS-TB) and drug-resistant (DR-TB) patients were recruited to examine immune profiles in these settings. Patients were followed up: at inclusion (T0), after two months of treatment (T1), and at the end of TB treatment (T2; 6 months for DS-TB patients, 9 to 24 months for DR-TB patients). The T1 timepoint was chosen because it marks the moment after which antibiotic treatment is reduced during clinical DS-TB management. For DR-TB monitoring, the same timepoint was used for consistency. Patients were on Directly Observed Treatment (DOT) and received treatment according to standard protocols (World Health Organization Geneva, 2019). Treatment regimens are detailed in Supp. Table 1.

### Whole blood stimulation and processing

Detailed whole blood collection and stimulation processes were described elsewhere (Chedid et al., 2021). Briefly, at every follow-up visit, 1mL of whole blood was drawn from the antecubital area of the arm and seeded directly into each QuantiFERON-TB Gold Plus (QFT-P, Qiagen) tube and incubated for 24 hours. Three stimulation conditions were used: NIL as unstimulated control; TB2 which tubes contain the *M. tuberculosis* antigenic peptides ESAT-6 (>15aa) and CFP-10 (8-13aa), which induce responses from CD4^+^ T lymphocytes (Petruccioli et al., 2016), and an undisclosed peptide pool inducing CD8^+^ T lymphocyte stimulation (Qiagen, 2017); rmsHBHA which tubes contain recombinant *M. tuberculosis* heparin-binding hemagglutinin generated in *M. smegmatis* at a final concentration of 5µg/mL and graciously provided by the Delogu laboratory, UNICATT, Rome, Italy (Delogu et al., 2011). After incubation, plasma separation, and red blood cell lysis with FACS lysing buffer (BD Biosciences) according to the manufacturer’s instructions, the resulting fixed white blood cells pellets were stored at −80°C. Cryopreserved samples were air-shipped in dry ice with freezing controls to the Mérieux Foundation Emerging Pathogens Laboratory in Lyon, France (International Center for Infectiology Research, INSERM U1111).

### Experimental procedure for mass cytometry

#### Sample preparation

Cryopreserved cells were thawed and resuspended in phosphate buffer saline (PBS) to a concentration of 3.5×10^6^cells/mL. Between 1 and 1.5×10^6^ cells from each sample were aliquoted for staining. Cells were incubated 10 minutes with FcR Blocking Reagent (6µL/10^6^ cells; Miltenyi Biotec) and heparin sodium salt reconstituted in Millipore water (36µg/10^6^ cells; Sigma-Aldrich) to reduce nonspecific staining (Rahman et al., 2016).

#### Panel design

A 29-marker panel of metal-labeled antibodies was used. All antibodies were obtained from Fluidigm (Supp. Table 8). Briefly, the panel contained 28 T-cell oriented surface markers (lineage markers, chemokine receptors, activation markers, and exhaustion markers) and one intracellular target (perforin).

#### Experimental design and barcoding

As the study followed a longitudinal design, samples from a same patient were acquired in the same barcoded batch of 3 timepoints and 3 stimulation conditions to reduce experimental variation. Palladium barcoding (Mei et al., 2015) (Cell-ID 20-Plex, Fluidigm) was performed according to the manufacturer’s instructions for simultaneous staining and data acquisition. For each barcoding run, 18 patient T-cell samples were stained with unique combinations of intracellular palladium isotopes (Figure 1). Patient batches were processed in a random order and investigators were blinded to patient sputum culture results during data collection.

#### Staining procedure

Extracellular staining was performed on pooled barcoded cells in Maxpar cell staining buffer (Fluidigm) for 30 minutes at room temperature. Intracellular staining (perforin) was performed in Maxpar Perm-S Buffer (Fluidigm) for 30 minutes at room temperature. Stained cells were then incubated for 10 minutes in 1.6% formaldehyde (FA) freshly prepared from 16% stock FA (Sigma-Aldrich). DNA staining was performed by overnight incubation at 4°C in 2mL of 125nM Cell-ID Iridium intercalator solution (Fluidigm). Cells were then washed, pelleted, and kept at 4°C until acquisition.

#### Data acquisition

Samples were analyzed on a CyTOF2 mass cytometer upgraded to Helios (Fluidigm) hosted by the AniRA cytometry facility (Structure Fédérative de Recherche Lyon Gerland, INSERM U1111, Lyon, France). Samples were filtered twice through a 50µm nylon mesh and resuspended in EQ™ Four Element Calibration Beads (Fluidigm) diluted to 0.5X in Maxpar ultra-pure water (Fluidigm), to reach an acquisition rate of 150-200 events per second (0.5 x 10^6^ cells/mL). Data were collected using the on-board Fluidigm software.

### Data analysis

All data analyses were performed in RStudio (version 1.3.1073 with R version 4.0.3) and FlowJo (version 10.7.1).

#### Data cleaning and preliminary manual gating

Signal normalization, concatenation, debarcoding, and conversion into Flow Cytometry Standard (FCS) 3.0 format were performed using the Helios Software (Fluidigm). Debarcoded files were imported into FlowJo and arcsinh-transformed (cofactor = 5). Gaussian parameters of the Helios system were used for doublet exclusion (Leipold et al., 2015), then ^191^Ir^+ 193^Ir^+^ single events were manually isolated, and debris (CD45^-^ events) and calibration beads (^140^Ce^+^ events) were excluded). A preliminary manual gating analysis was then performed on CD45^+^ single events (Supp. Figure 8) to verify that the proportions of the main white blood cell subpopulations in biobanked samples were consistent with the expected proportions, and sufficient for downstream analysis. Samples with less than 1,000 CD3^+^ events, and batches with missing samples from a given timepoint were removed from the analysis to preserve a matched sample design. The exact number of available files per patient and per stimulation condition is provided in Supp. Table 1.

#### Workflow for unsupervised analyses

CD3^+^ single events were down-sampled to ensure equal contribution of each sample, exported into separate Comma Separated Value (.csv) files, and uploaded into R software (version 4.0.3). Panel markers were defined as either lineage or functional markers for use as clustering channels in downstream analyses (Supp. Table 9). Lineage-defining markers included canonical surface markers such as CD4 which display a theoretically stable expression. Functional markers included markers of activation (*e.g.* CD69), proliferation (CD38), maturation (CD27), or migration (CCR7).

#### Dimension reduction, automated clustering, and phenotyping

After file concatenation, dimension reduction was performed with UMAP (Uniform Manifold Approximation and Projection; version 3.1) (Becht et al., 2019). UMAPs were created in R using the package Spectre (Ashhurst et al., 2020). Unsupervised clustering was performed using FlowSOM (Van Gassen et al., 2015) (version 2.7). FlowSOM meta-cluster phenotyping was assessed by visualizing the surface expression of lineage markers in each FlowSOM cluster (CD4, CD8, TCRgd, TCRVa7.2, CD56, CD25, IL7Ra, CD26, and CD161) on a heatmap and performing hierarchical clustering. Marker expression heatmaps were obtained in R using Spectre by plotting normalized, median arcsinh-transformed mass signals. Biological consistency of FlowSOM meta-clusters with the main expected T-cell subpopulations (Supp. Table 3) was controlled, and manual reassignment of clusters which were in inconsistent meta-clusters was then performed when necessary (Supp. Figure 9). Meta-clusters with an abundance <1% of all events were pooled with the most phenotypically similar meta-cluster. Then, the proportion of corrected FlowSOM meta-clusters in each node on the initial FlowSOM minimum spanning tree was visualized to control reassignment consistency (Quintelier et al., 2021).

#### Statistical analysis

The proportion (percent of CD3^+^) of each FlowSOM cluster was calculated. For all statistical analyses, exact p-values, test statistics and/or estimates of effect size are provided either in the figure legend or in indicated Supplementary Tables. Normality was assessed using the Shapiro-Wilk test. The evolution of cluster proportions over time corresponded to repeated measures of non-normal, non-independent continuous variables, and was analyzed in matched samples using the two-sided Friedman rank sum test with the Wilcoxon–Nemenyi–McDonald-Thompson post-hoc test (Pereira et al., 2015). Independent, non-normal continuous variables were analyzed with the two-sided Mann–Whitney U test or the Kruskal–Wallis test with Dunn’s Kruskal–Wallis Multiple Comparisons *post-hoc* test (Dunn, 1964) when more than two categories were compared. For discovery of clusters with significantly different abundance between slow and fast converters, conservative corrections for multiple comparisons (*e.g.* Benjamini-Hochberg (Yoav Benjamini and Yosef Hochberg, 1995)) were not used in order to minimize type II errors. Instead, all p-values were computed for each timepoint, and the p-value corresponding to the null hypothesis being rejected in 5% of all comparisons was used as the significance threshold instead of 0.05 (Althouse, 2016). This novel significance threshold enabled to control type I error while maintaining an exploratory approach; its value was always inferior to 0.05 and is reported in the corresponding figure captions.

## Supplemental Materials

Supp. Figure 1 is a flowchart of patient inclusions. Supp. Figure. 2 shows the impact of *in vitro Mtb* antigen whole blood stimulation on surface marker expression in the main T-cell compartments. Supp. Figure. 3 summarizes the frequencies of the main peripheral T-cell subpopulations throughout anti-TB treatment. Supp. Figures 4 to 6 quantify and illustrate how patients with slow microbiological culture conversion show decreased CD8^+^ and enriched CD4^+^ naïve peripheral T-cell subsets during treatment. Supp. Figure 7 reports PCA data characterizing the variance between fast and slow responders within all *Mtb*-stimulated CD3^+^ T-cells. Supp. Figure 8 and 9 relate to the Methods section, and show the main CD45^+^ non-granulocyte whole blood subpopulations, a T-cell oriented gating strategy, as well as the methods used to control automated FlowSOM metaclustering. Regarding tables, Supp. Table 1 summarizes the sociodemographic and clinical characteristics of the cohort. Supp. Tables 2 through 7 list the exact p-values and test statistics for all analyses presented throughout the main manuscript. Supp. Table 8 lists the mass cytometry panel components. Supp. Table 9 shows how clustering channels were defined in relation to expected cell subpopulations for dimension reduction and automated clustering of CD3^+^ T-cells.

## Author contributions

FA and JH are the principal investigators and initiated the project together with DG, NT, and SBa. Samples were collected by EK, NT, MU, and SBi. CC and TA designed and optimized the mass cytometry protocol. CC performed all experiments and analyses. CC and FA wrote the manuscript. All authors contributed to the article and approved the submitted version.

## Acknowledgements

We would like to thank the patients participating in our study, as well as the healthcare staff and laboratory collaborators in each study site.

## Data availability

The datasets generated and used in this study are available from the corresponding author upon reasonable request, excluding confidential patient information.

