## Supplemental tables and figures for "In-depth immunophenotyping with mass cytometry during TB treatment reveals non-canonical T-cell subsets associated with sputum culture conversion"

### Supplementary Information

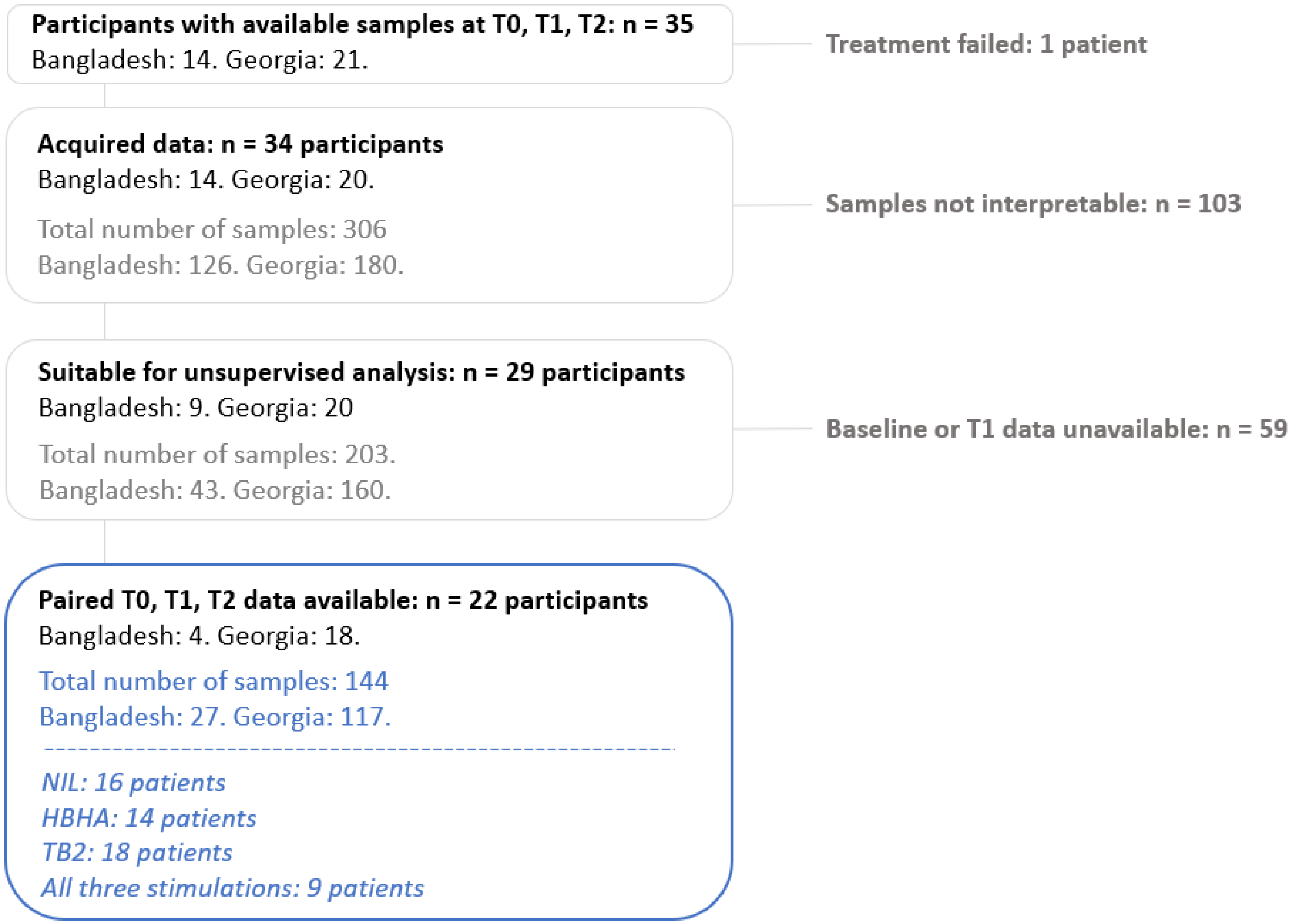


**Supp. Figure 1. Flowchart of patient inclusions.**

A table listing the number of cell samples available for each stimulation condition and each patient individually is provided in Supp Table 1.

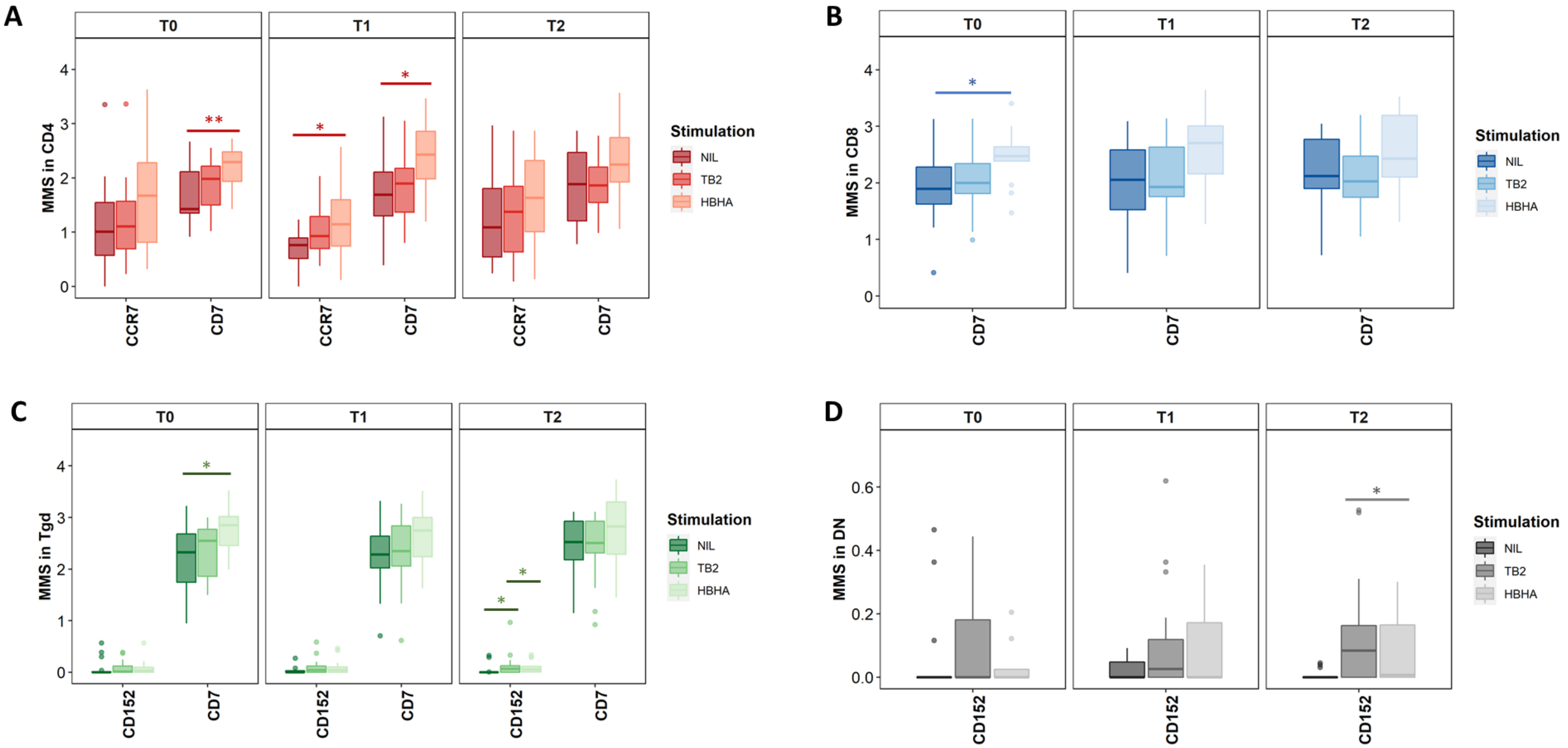


**Supp. Figure. 2. Impact of *in vitro* whole blood stimulation with *Mtb* antigens on surface marker expression in the main T-cell subpopulations.** The surface expression of all panel markers was compared between the three stimulation conditions (unstimulated (NIL), TB2, and rmsHBHA) in CD4$^+$ (**A**), CD8$^+$ (**B**), gamma-delta (Tgd; **C**), or double negative (DN) T-cells (**D**). MMS: median mass signal. Only the markers for which a significant difference was observed were represented. Statistical analysis: two-sided Kruskal-Wallis test with Dunn's Kruskal–Wallis Multiple Comparisons post-hoc at T0, T1, and T2. *: $p<0.05$. **: $p<0.01$. Number of data points per timepoint for all panels: NIL: n = 16. TB2: n = 18. HBHA: n = 14. Exact p-values and test statistics are available in Supp. Table 2.

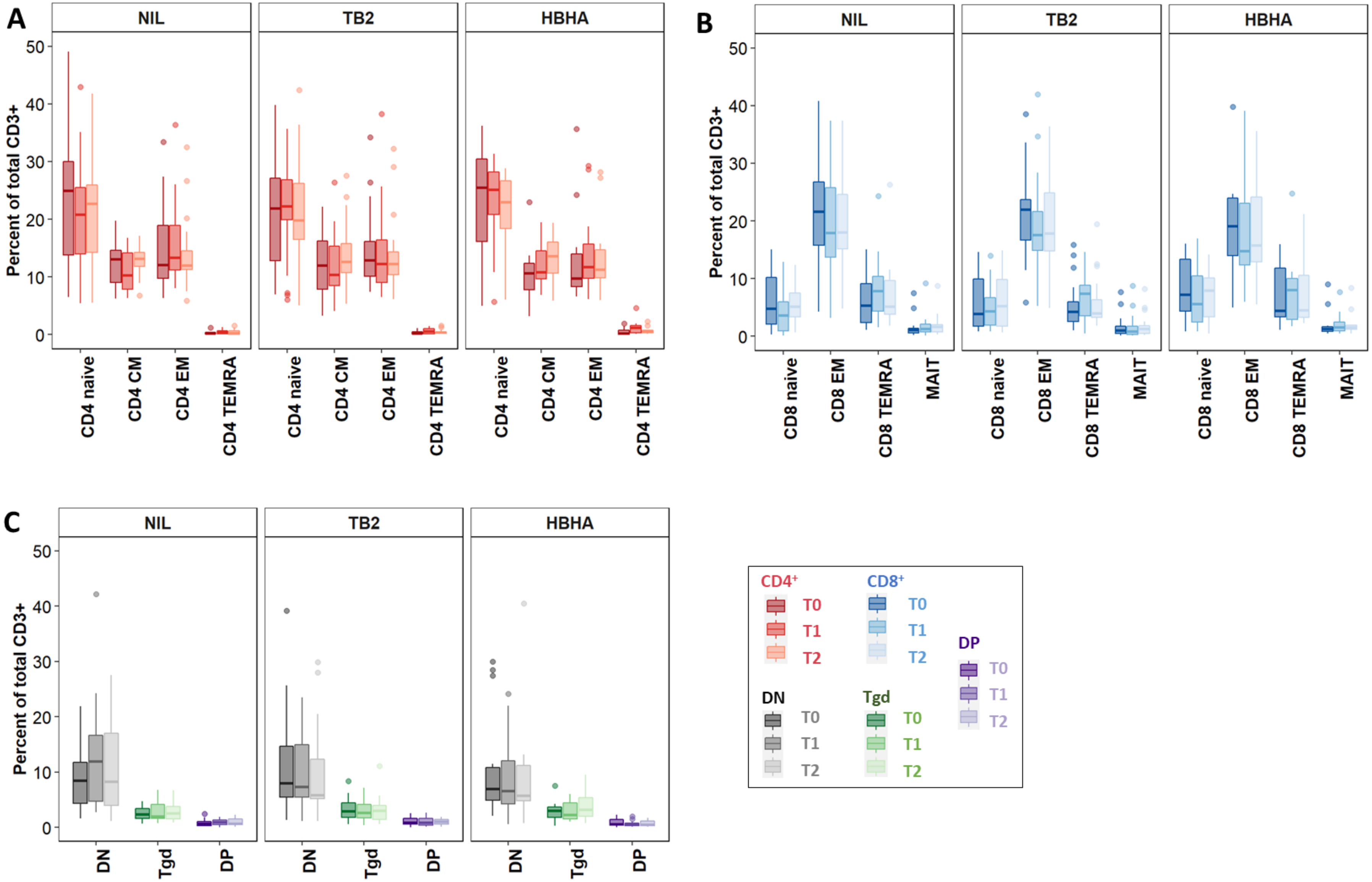
A
NIL
TB2
HBHA
Percent of total CD3+
0
10
20
30
40
50
CD4 naive
CD4 CM
CD4 EM
CD4 TEMRA
B
NIL
TB2
HBHA
Percent of total CD3+
CD8 naive
CD8 EM
CD8 TEMRA
MAIT
C
NIL
TB2
HBHA
Percent of total CD3+
DN
Tgd
DP
CD4+
T0
T1
T2
CD8+
T0
T1
T2
DP
T0
T1
T2
DN
T0
T1
T2
Tgd
T0
T1
T2

**Supp. Figure. 3. Frequencies of the main peripheral T-cell subpopulations throughout anti-TB treatment.**

Evolution of the frequency of canonical T-cell subsets identified through FlowSOM meta-clustering and corresponding respectively to $CD4^+$ phenotypes (**A**), $CD8^+$ phenotypes (**B**), or other cell subsets (**C**). Number of data points per timepoint for all panels: NIL: n = 16. TB2: n = 18. HBHA: n = 14. Data are given as median + interquartile range. Abbreviations: CM: central memory. DN: double-negative $CD4^-CD8^-$. DP: double-positive $CD4^+$ $CD8^+$. EM: effector memory. HBHA: recombinant *M. tuberculosis* heparin-binding hemagglutinin. MAIT: mucosal associated invariant T-cells. NIL: unstained control. TB2: *M. tuberculosis* antigenic peptide pool. Tgd: gamma delta T-cells. Treg: T-regulators. TEMRA: terminally differentiated effectors re-expressing CD45RA. No statistically significant differences were detected (pairwise comparisons between non-sindependent observations at T0, T1, and T2: two-sided Friedman rank sum test and Wilcoxon-Nemenyi-Thompson post-hoc for pairwise comparisons between non-independent observations at T0, T1, and T2).

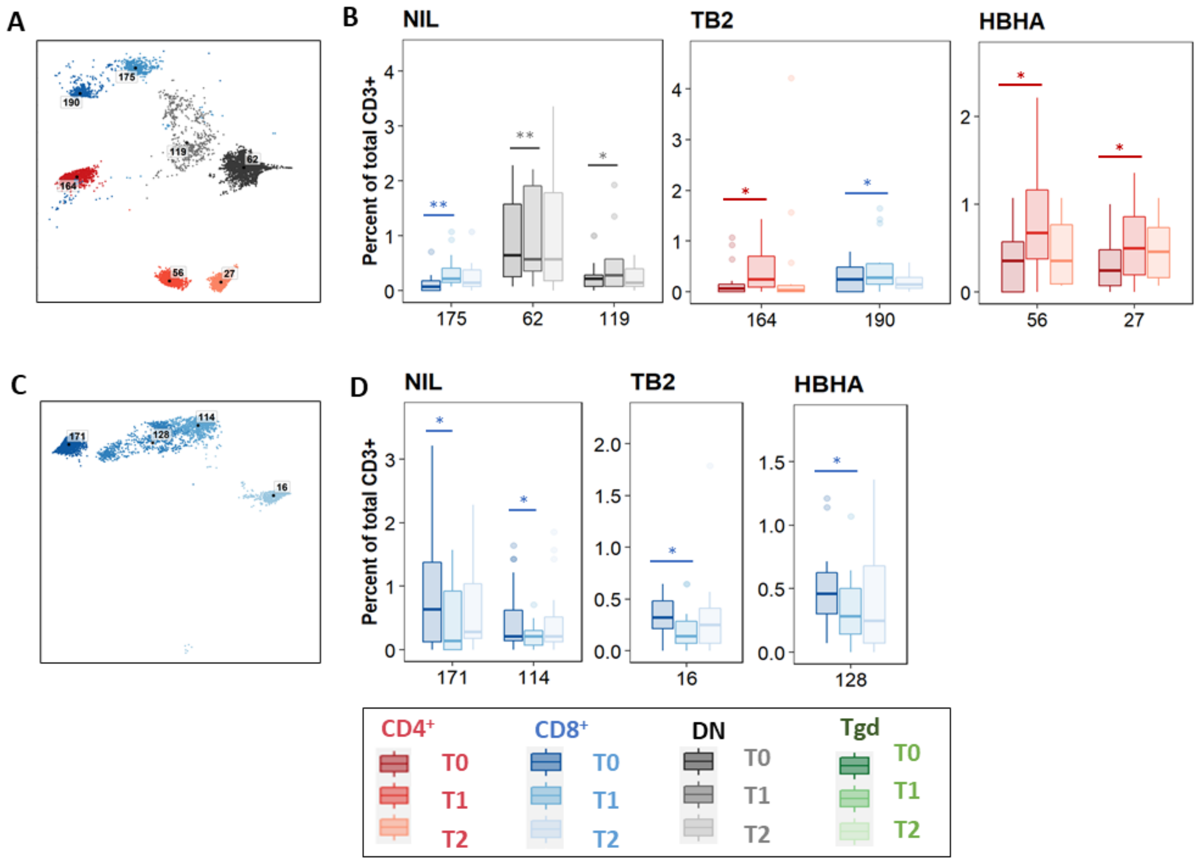


**Supp Figure 4. Significant abundance changes in non-canonical T-cell subsets after the intensive phase of treatment.** The evolution of FlowSOM cluster abundance was analyzed over time in unstimulated or *Mtb*-stimulated samples (TB2 or rmsHBHA), and only the clusters within which significant abundance changes were detected were displayed. $CD4^+$ clusters were represented in red, $CD8^+$ clusters in blue, γδ T-cell clusters in green, and $CD4^-$ $CD8^-$ clusters in grey. Number of matched data points per timepoint for all panels: NIL: n = 16. TB2: n = 18. rmsHBHA: n = 14. Data are given as median + interquartile range.

**A and B.** Significantly increased clusters at the end of the intensive phase of treatment (T1) compared to treatment initiation (T0). Clusters within which a significant increase was detected between T0 and T1 were first visualized on the reference UMAP (**A**). Cluster abundance quantification was then was performed in unstimulated, TB2-stimulated or rmsHBHA-stimulated samples (**B**).

**C and D**. Significantly decreased clusters at the end of the intensive phase of treatment (T1) compared to treatment initiation (T0). Mapping **(C)** and abundance quantification of clusters which decreased between T0 and T1 in unstimulated, TB2-stimulated, or rmsHBHA-stimulated samples (**D**).

<u>Statistical analysis</u>: two-sided Friedman rank sum test and Wilcoxon-Nemenyi-Thompson post-hoc for pairwise comparisons between non-independent observations at T0, T1, and T2. *: p<0.05. **: p<0.01. ***: p<0.001. Exact p-values and test statistics are available in Supp. Table 3 (associated Excel file).

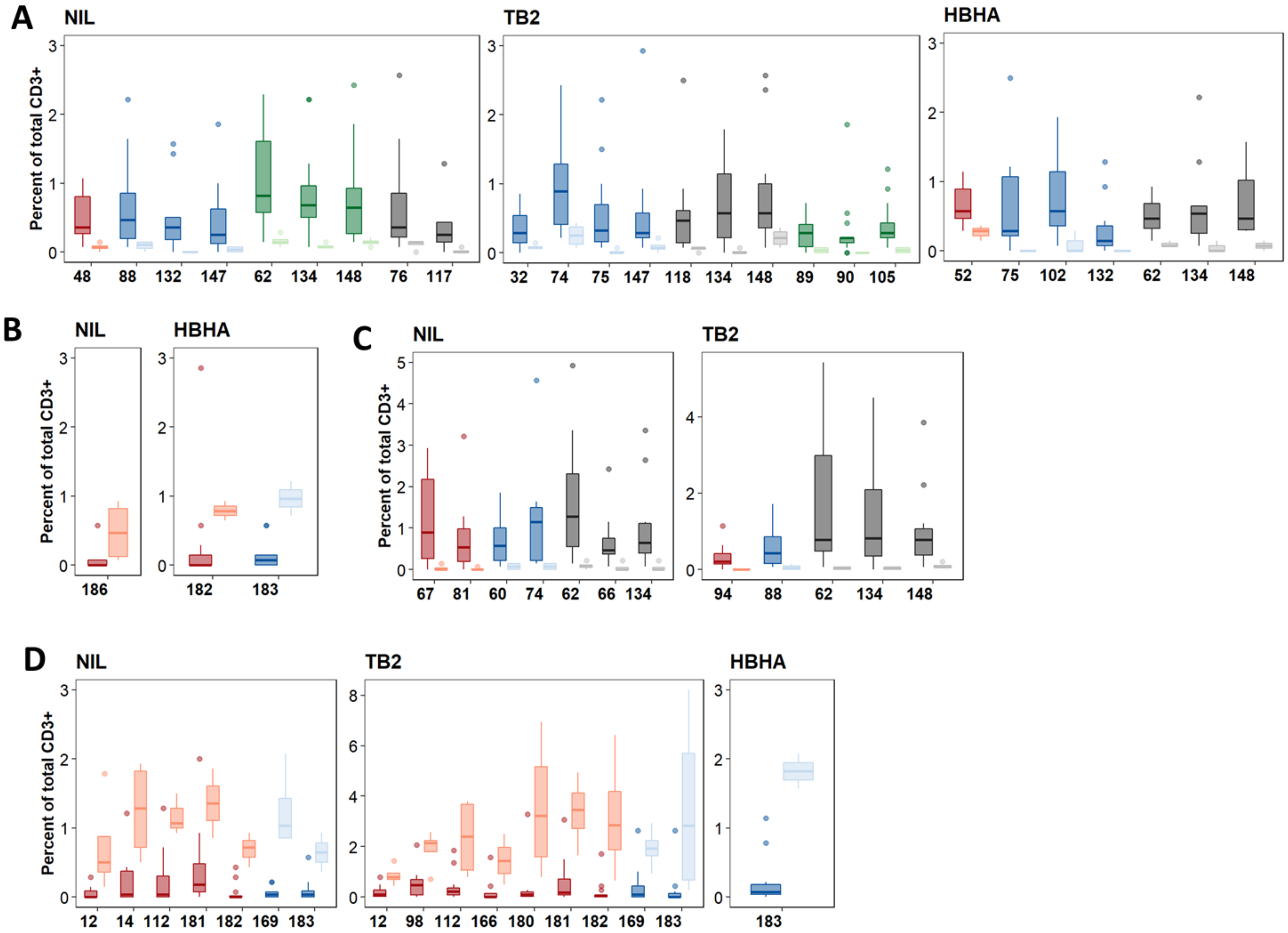

A
NIL
TB2
HBHA
Percent of total CD3+
48 88 132 147 62 134 148 76 117
32 74 75 147 118 134 148 89 90 105
52 75 102 132 62 134 148
B
NIL
HBHA
186
182 183
C
NIL
TB2
67 81 60 74 62 66 134
94 88 62 134 148
D
NIL
TB2
HBHA
12 14 112 181 182 169 183
12 98 112 166 180 181 182 169 183
183

**Supp. Figure. 5. Patients with slow microbiological culture conversion show decreased CD8$^+$ and γδ and enriched CD4$^+$ naïve peripheral T-cell subsets during treatment.**

Clusters with differential abundance between patients with positive mycobacterial cultures at T1 (slow converters, n = 4) or with negative cultures at T1 (fast converters, n = 18). Data are shown as median + interquartile range.

**A and B. At treatment initiation (T0).** Clusters significantly decreased (**A**) and increased (**B**) in slow converters compared to fast converters.

**C and D. At treatment completion (T2).** Clusters significantly decreased (**C**) and increased (**D**) in slow converters compared to fast converters.

CD4$^+$ clusters are represented in red, CD8$^+$ clusters in blue, Tgd clusters in green, and DN clusters in grey. The lighter shade of each color code corresponds to data from the slow converters. Statistical analysis: two-sided Mann-Whitney U-test. For all represented clusters: P<0.025 at T0; P<0.013 at T2. Significance stars were not displayed for readability. Exact p-values and test statistics are available in Supp. Table 6 (associated Excel file).

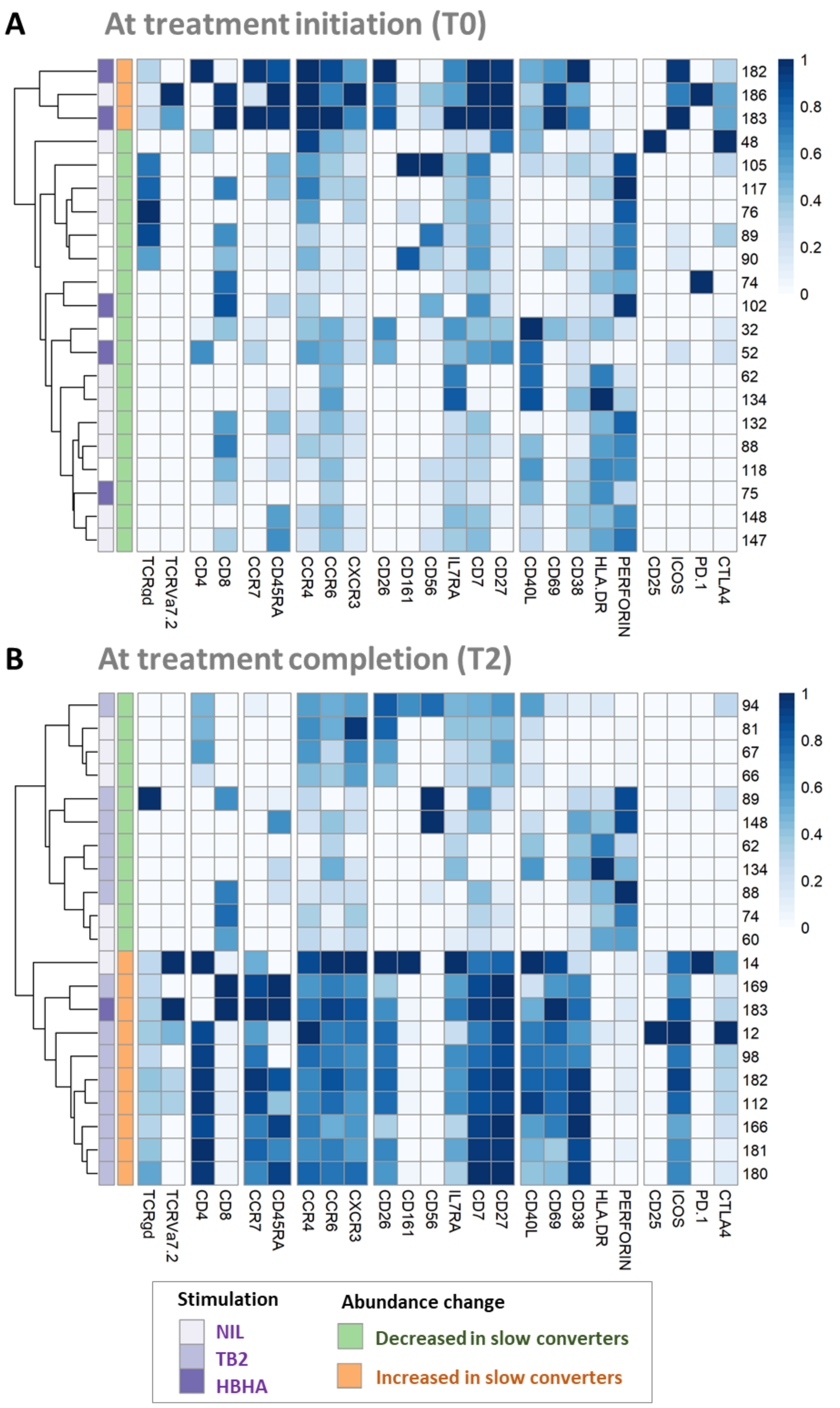
A
At treatment initiation (T0)
182
186
183
48
105
117
76
89
90
74
102
32
52
62
134
132
88
118
75
148
147
TCRgd
TCRVa7.2
CD4
CD8
CCR7
CD45RA
CCR4
CCR6
CXCR3
CD26
CD161
CD56
IL7RA
CD7
CD27
CD40L
CD69
CD38
HLA.DR
PERFORIN
CD25
ICOS
PD.1
CTLA4
1
0.8
0.6
0.4
0.2
0
B
At treatment completion (T2)
94
81
67
66
89
148
62
134
88
74
60
14
169
183
12
98
182
112
166
181
180
Stimulation
NIL
TB2
HBHA
Abundance change
Decreased in slow converters
Increased in slow converters

**Supp. Figure 6. Patients with slow microbiological culture conversion show decreased cytotoxic CD8$^{+}$ and γδ and enriched CD4$^{+}$ naïve T-cell subsets before treatment initiation and after treatment completion compared to fast converters.** Fast converters (n = 18) were defined as patients with permanently negative *M. tuberculosis* culture after the intensive phase of treatment (T1), whereas slow converters (n = 4) were defined as patients with persistently positive cultures at T1. The abundance of all FlowSOM clusters at baseline was compared between fast and slow converters. Only clusters within which significant differences were detected were represented (T0: $p<0.026$. T2: $p<0.013$; two-sided Mann-Whitney U test; see Supp. Figure 5).

**A. Before treatment initiation (T0).** Clusters which were significantly decreased (green) or increased (orange) at T0 in slow converters compared to fast converters were represented. Normalized, arcsinh-transformed mean marker expression levels were visualized). Each line represents one cluster. Scales indicate normalized mass signal intensity.

**B. After treatment completion (T2).** Clusters which were significantly increased or decreased at T2 in slow converters compared to fast converters were represented and marker expression levels were visualized. All patients achieved microbiological cure at T2.

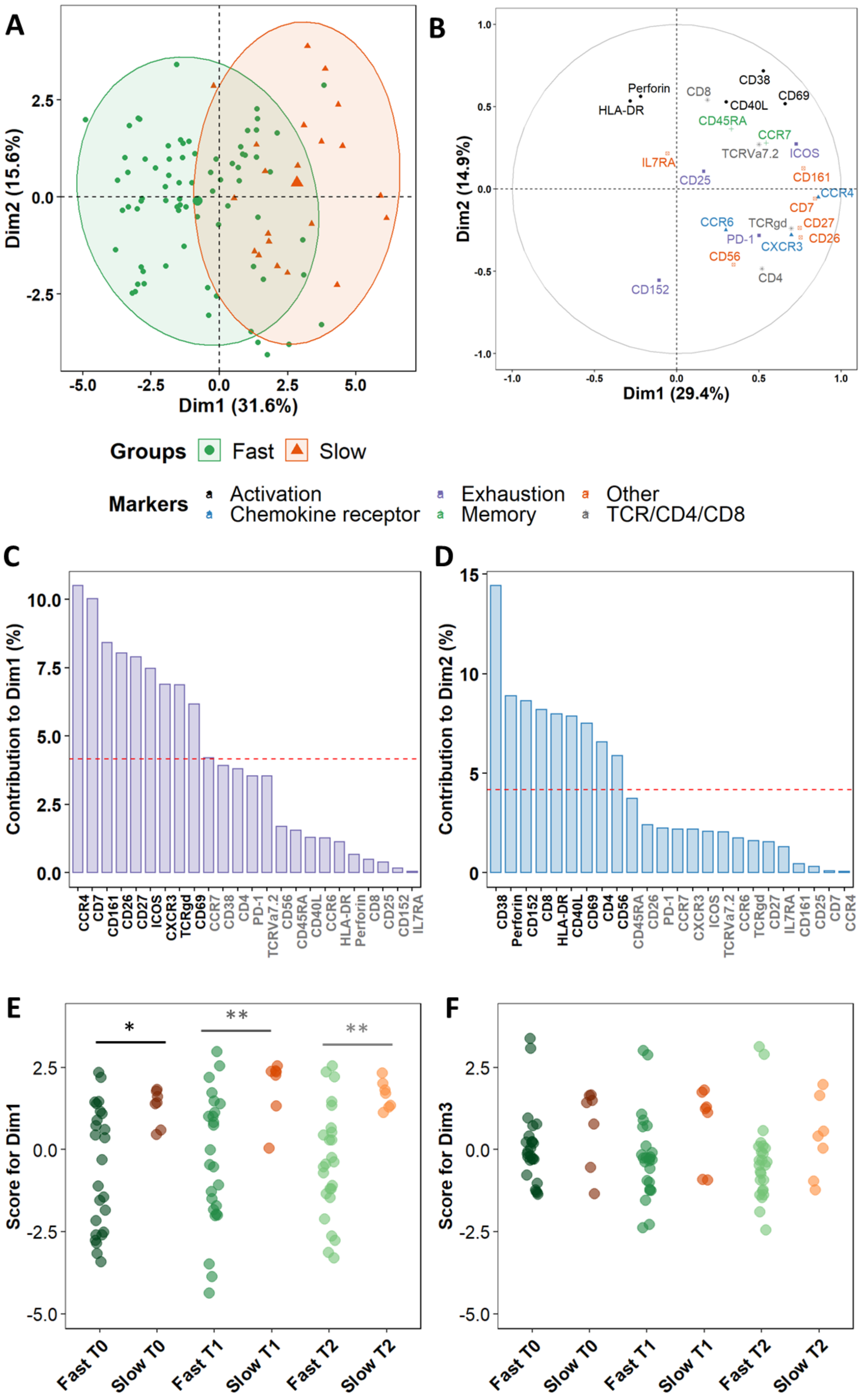
A
B
C
D
E
F
Dim1 (31.6%)
Dim2 (15.6%)
Dim1 (29.4%)
Dim2 (14.9%)
Groups
Fast
Slow
Markers
Activation
Chemokine receptor
Exhaustion
Memory
Other
TCR/CD4/CD8
Contribution to Dim1 (%)
Contribution to Dim2 (%)
Score for Dim1
Score for Dim3
Fast T0
Slow T0
Fast T1
Slow T1
Fast T2
Slow T2

**Supp. Figure 7. Variance between fast and slow responders within all *Mtb*-stimulated $CD3^+$ T-cells.**

Principal Component Analysis (PCA) was performed on marker expression data within all $CD3^+$ T-cells from 96 *Mtb*-stimulated samples matched at T0, T1, and T2 (TB2: 54 samples; rmsHBHA: 42 samples).

**A.** Explanation of the variance between fast converters (25 samples at each timepoint) and slow converters (7 samples at each timepoint). Each dot represents one patient. The color code represents the culture conversion group. Axes represent the principal components 1 (Dimension 1, Dim1) and 2 (Dim2) and percentages indicate their contribution to the total observed variance. Axis values represent individual PCA scores. Concentration ellipses correspond to 90% data coverage.

**B.** Contribution of cellular markers to the variance described by Dim1 and Dim2. Axis values represent marker PCA scores. The color code represents broad marker functions.

**C and D.** Quantification of panel B. for Dim1 (**C**) and Dim2 (**D**). Contributions of each marker are expressed as a percentage of the dimensions. The red dashed line corresponds to the expected reference value if each marker contributed uniformly to the variance. Markers indicated in gray are below this reference value.

**E and F.** Distribution of individual PCA score values according to the culture conversion group and to the timepoint, for Dim1 (**E**) and Dim2 (**F**). Data were compared with the two-sided Wilcoxon Rank Sum Test. *: $p<0.05$; **: $p<0.01$. Exact p-values and test statistics are available in Supp. Table 7.

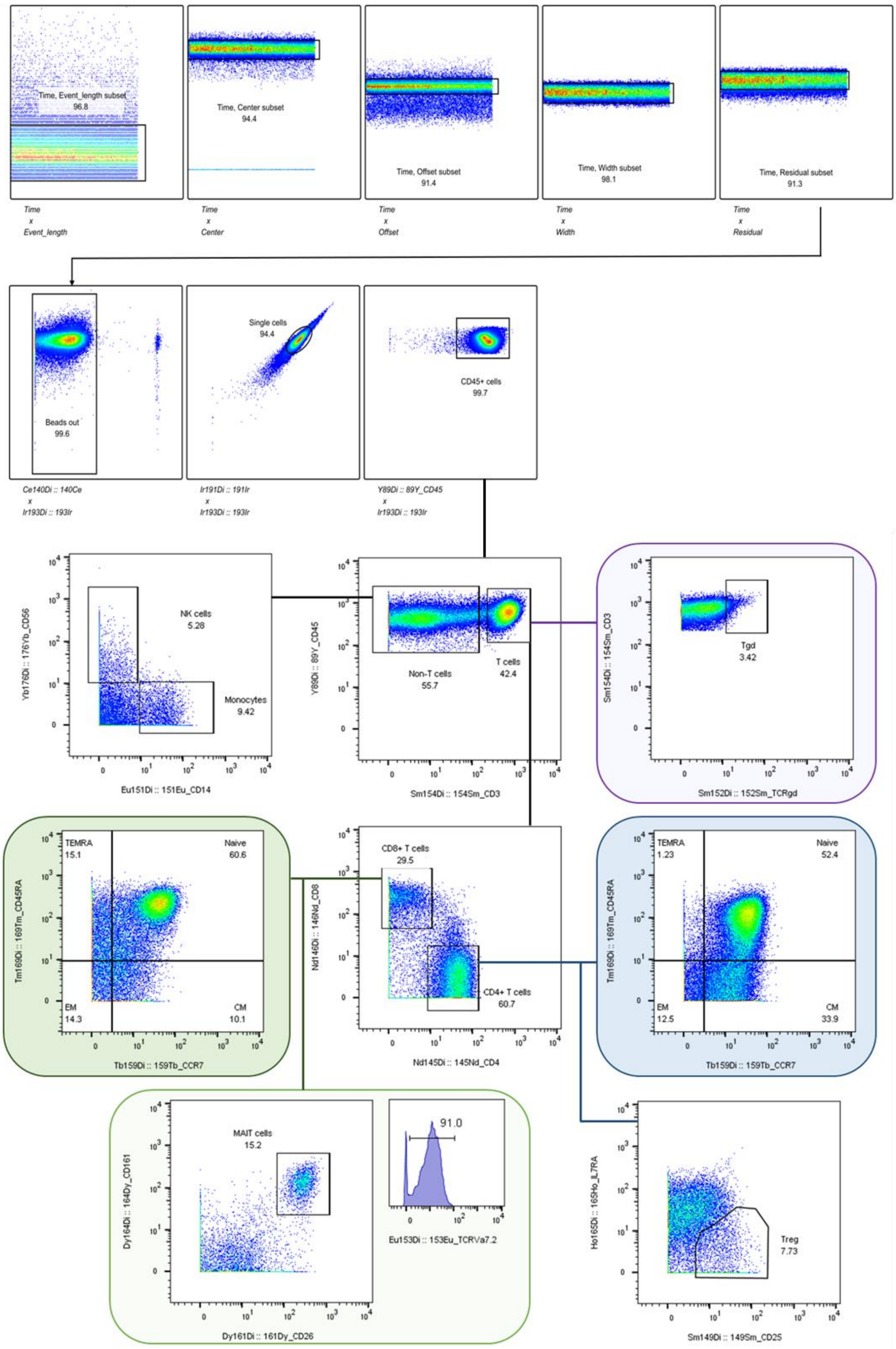


**Supp. Figure 8. Main CD45+ non-granulocyte whole blood subpopulations and T-cell oriented gating strategy.** CM: central memory. EM: effector memory MAIT: mucosal-associated invariant T-cells. NK: natural killers. TEMRA: terminally differentiated effectors re-expressing CD45RA. Tgd: gamma delta T-cells. Treg: T regulators.

A

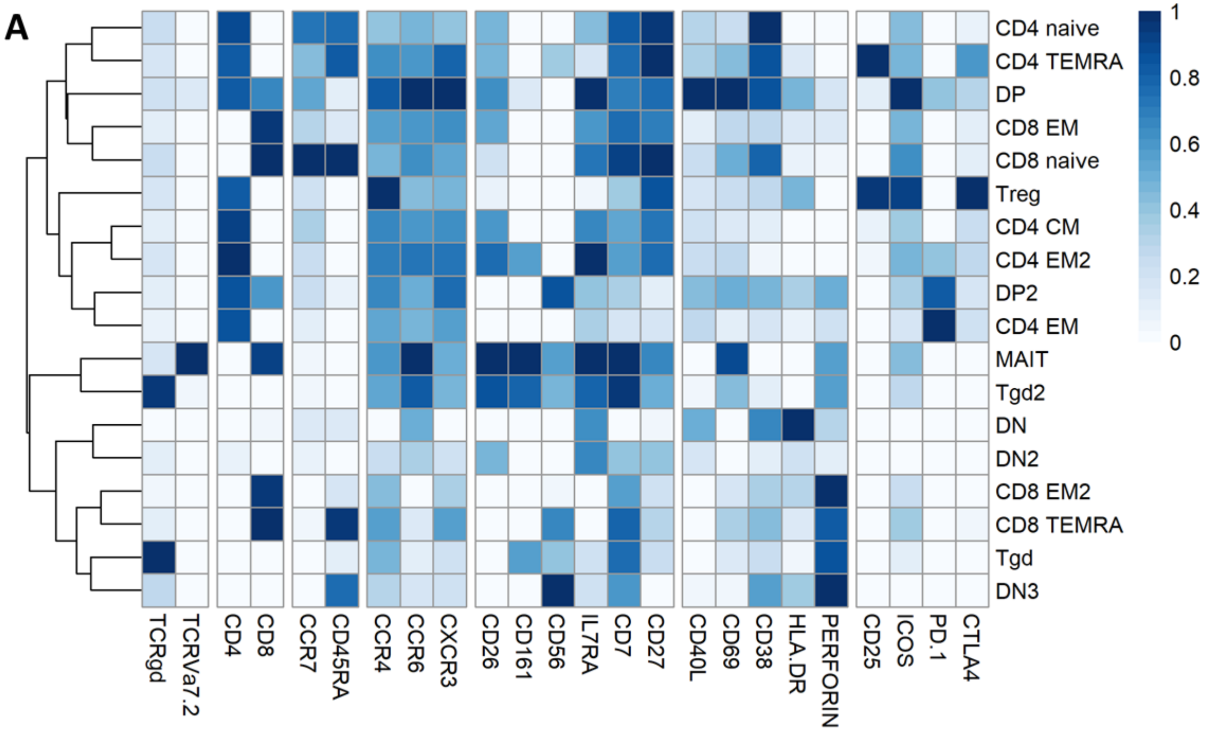


B

| Initial metacluster | Merged with |
|---|---|
| CD4 EM2 | CD4 EM |
| Tgd2 | Tgd |
| CD8 EM2 | CD8 EM |
| DN2 | DN |
| DN3 | |
| DP2 | DP |
| DN | CD8 EM |

**Supp. Figure 9. Control of automated FlowSOM metaclustering.**

**A.** Expression of selected lineage markers in 18 automatically detected FlowSOM meta-clusters within total $CD3^{+}$ events. A higher number of meta-clusters than expected was chosen in order to detect all expected cell subpopulations (see Supp.Table 9). **B.** List of meta-clusters which were reassigned to other phenotypically similar meta-clusters. <u>Abbreviations:</u> CM: central memory. DN: double-negative $CD4^{-}CD8^{-}$. DN: double-positive $CD4^{+}$ $CD8^{+}$. EM: effector memory. HBHA: recombinant *M. tuberculosis* heparin-binding hemagglutinin. MAIT: mucosal associated invariant T-cells. NIL: unstained control. TB2: *M. tuberculosis* antigenic peptide pool. Tgd: gamma delta T-cells. Treg: T-regulators. TEMRA: terminally differentiated effectors re-expressing CD45RA.

**Supplementary Table 1. Sociodemographic and clinical characteristics of the cohort.**

| | Nb of cell samples | | | Drug susceptibility | | |
|---|---|---|---|---|---|---|
| **ID** | **NIL** | **TB2** | **HBHA** | **Phenotypic drug susceptibility** | **Treatment regimen** | **Country** |
| 01DSGE | 3 | 3 | 3 | DS | 2HRZE/4HRE | GEO |
| 02DSGE | 3 | 0 | 3 | DS | 2HRZE/4HRE | GEO |
| 03DSGE | 3 | 3 | 3 | DS | 2HRZE/4HR | GEO |
| 04DSGE | 3 | 0 | 0 | DS | 2HRZE/4HR | GEO |
| 05DSGE | 3 | 3 | 3 | DS | 2HRZE/4HR | GEO |
| 06DSB | 3 | 3 | 0 | DS | 2HRZE/4HR | BD |
| 06DSGE | 3 | 3 | 0 | DS | 2HRZE/4HRE | GEO |
| 07DSGE | 3 | 3 | 3 | DS | 2HRZE/4HRE | GEO |
| 08DSB | 3 | 3 | 0 | DS | 2HRZE/4HR | BD |
| 08DSGE | 3 | 3 | 3 | DS | 2HRZE/4HRE | GEO |
| 21DSB | 0 | 3 | 3 | DS | 2HRZE/4HR | BD |
| 01DRGE | 0 | 3 | 0 | MDR | H+Z+Km+Lfx+cfz+Bdq / Z+Lfx+cfz+Bdq | GEO |
| 02DRGE | 3 | 3 | 3 | MDR | Cm+Lfx+Pto+Cs+Cfz+H / Cm+Lfx+Cs+Bdq+Lzd / Lfx+Cs+Bdq+Lzd | GEO |
| 03DRGE | 3 | 0 | 3 | Pre-XDR | (Z)+Lfx+Pto+Cfz+Bdq+Lzd / Cs+Cfz+Bdq+Lzd | GEO |
| 04DRGE | 0 | 3 | 0 | Pre-XDR | Bdq+Cfz+Lfx+H+Pto+E+Z / Bdq+cfz+Lzd+H+E / Bdq+Cfz+Lfx+H+Z / Bdq+Cfz+Lfx+E+Z | GEO |
| 05DRGE | 3 | 3 | 0 | Pre-XDR | H+Z+Km+Lfx+Cfz+Bdq / Z+Lfx+Cfz+Bdq | GEO |
| 06DRGE | 0 | 3 | 3 | RR | H+E+Z+Km+Mfx+Pto+Cfz / Mfx+E+Z+cfz | GEO |
| 07DRGE | 3 | 3 | 3 | XDR | Bdq+Lzd+cfz+Cs | GEO |
| 09DRB | 3 | 3 | 3 | MDR | 4-6 Km-Mfx-Pto-Cfz-Z-HHigh dose-E/5-6 Mfx-Cfz-Z-E3 | BD |
| 09DRGE | 0 | 0 | 3 | MDR | H+Z+Km+Lfx+Cfz+Bdq / Z+Lfx+Cfz+Bdq | GEO |
| 10DRGE | 0 | 3 | 0 | MDR | H+Z+Km+Lfx+Cfz+Bdq / Z+Lfx+Cfz+Bdq | GEO |
| 14DRGE | 3 | 3 | 3 | Pre-XDR | H+Z+Km+Lfx+Cfz+Bdq / Z+Lfx+Cfz+Bdq | GEO |
| ***Nb of patients:*** | *16* | *18* | *14* | | | |
| **Number of samples:** | 48 | 54 | 42 | | | |
| **Total number of samples:** | **144** | | | | | |

Footnotes: Three samples per patient were collected in each stimulation condition, corresponding to each timepoint: T0: baseline. T1: T0 + 2 months. T2: end of treatment.Samples with < 1,000 $CD3^{+}$ events, and batches with missing samples from a given timepoint were removed from the analysis. Abbreviations: BD: Bangladesh. GEO: Georgia. DS: drug susceptible. MDR: multi-drug resistant. RR: rifampicin resistant. XDR: extensively drug resistant.

Abbreviations for anti-TB drugs: E: Ethambutol. H: Isoniazid. R: Rifampicin. S: Streptomycin. Z: Pyrazinamide. Bdq: Bedaquiline. Cfz: Clofazimine. Cs: Cycloserine. Km: Kanamycin. Lfx: Levofloxacin. Lzd: Linezolid. Mfx: Moxifloxacin. Pto: Prothionamide. Numbers indicate months of treatment when the information was available

**Supplementary Table 1 - continued**

| Baseline identification | | | | | | | | | | Microbiological evolution | | | | | | |
|---|---|---|---|---|---|---|---|---|---|---|---|---|---|---|---|---|
| ID | Gender | Age | Occupation | HIV | Diabetes | BCG | QFT-P | GeneXpert | GeneXpert RIFR | *Mtb* culture | | | AFB | | | Treatment response group |
| | | | | | | | | | | T0 | T1 | T2 | T0 | T1 | T2 | |
| 01DSGE | M | 28 | Small business | n | n | | + | + | - | + | - | - | scanty | scanty | - | Fast |
| 02DSGE | M | 26 | Student | n | n | y | + | + | - | + | - | - | - | - | - | Fast |
| 03DSGE | M | 37 | Small business | n | n | y | + | + | - | + | - | - | - | - | - | Fast |
| 04DSGE | M | 24 | Day Laborer | n | n | y | + | + | - | + | - | - | - | - | - | Fast |
| 05DSGE | M | 63 | Unemployed / Retired | n | n | | + | + | - | + | - | - | 2+ | 2+ | - | Fast |
| 06DSB | M | 16 | Day Labour | n | n | y | + | + | - | + | - | - | 2+ | scanty | - | Fast |
| 06DSGE | M | 58 | Unemployed / Retired | n | n | | - | + | - | + | + | - | 1+ | 1+ | scanty | Slow |
| 07DSGE | M | 50 | Unemployed / Retired | n | n | | + | + | - | + | + | - | scanty | scanty | - | Slow |
| 08DSB | M | 60 | Small business | n | n | y | - | + | - | + | - | - | 1+ | - | - | Fast |
| 08DSGE | F | 29 | Small business | n | n | y | + | + | - | + | + | - | scanty | scanty | - | Slow |
| 21DSB | F | 20 | Student | n | n | y | + | + | - | + | - | - | 2+ | - | - | Fast |
| 01DRGE | M | 42 | Farmer | n | n | | - | + | - | + | - | - | 1+ | - | - | Fast |
| 02DRGE | F | 34 | Housewife | n | n | y | - | + | + | + | + | - | - | - | - | Slow |
| 03DRGE | M | 26 | Business | n | n | y | + | + | - | + | - | - | 2+ | - | - | Fast |
| 04DRGE | M | 37 | Private Service | n | n | | + | + | - | + | - | - | - | 1+ | - | Fast |
| 05DRGE | M | 42 | Unemployed | n | n | | + | + | - | + | - | - | 3+ | scanty | - | Fast |
| 06DRGE | F | 33 | Unemployed | n | n | | + | + | + | + | - | - | 2+ | - | - | Fast |
| 07DRGE | F | 20 | | n | n | y | - | + | + | + | - | - | 2+ | - | - | Fast |
| 09DRB | F | 15 | Garments worker | n | n | y | - | + | + | + | - | - | 3+ | - | - | Fast |
| 09DRGE | F | 28 | | n | n | | + | + | - | + | - | - | 2+ | - | - | Fast |
| 10DRGE | F | 25 | | n | n | | - | + | + | + | - | - | - | - | - | Fast |
| 14DRGE | M | 31 | | n | n | | + | + | + | + | - | - | scanty | scanty | scanty | Fast |

Footnotes**:** BCG: Bacille Calmette-Guérin vaccination. QFT-P: QuantiFERON-TB Gold Plus. AFB: Acid Fast Bacilli detection (sputum smear microscopy). BMI: body mass index. *Mtb: Mycobacterium tuberculosis*. Throughout the table, "+" and "-" indicate positive or negative results to the indicated test. For AFB results, 1+, 2+, or 3+ quantify the amount of bacilli observed.

**Supplementary Table 1 - continued**

| Weight evolution | | | | | | | TB risk factors | | | | | |
|---|---|---|---|---|---|---|---|---|---|---|---|---|
| ID | Weight (kg) | | | BMI | | | Smoking | Alcohol | Intraveinous drug use | Prison | TB contact | Previous TB |
| | T0 | T1 | T2 | T0 | T1 | T2 | | | | | | |
| 01DSGE | 72 | 72 | 73 | 20.8 | 20.8 | 21.1 | n | n | n | n | n | n |
| 02DSGE | 59 | 61 | 64 | 20.1 | 20.8 | 21.8 | n | n | n | n | n | n |
| 03DSGE | 65 | 66 | 71 | 24.1 | 24.5 | 26.3 | y | n | n | n | n | n |
| 04DSGE | 59 | 60 | 64 | 19.9 | 20.2 | 21.6 | y | n | n | n | n | n |
| 05DSGE | 66.7 | 67 | 68 | 24.2 | 24.3 | 24.6 | n | n | n | n | n | n |
| 06DSB | 42.2 | 43.3 | 48.1 | 16.4 | 16.9 | 18.7 | n | n | y | n | n | n |
| 06DSGE | 64.2 | | | 18.3 | | | y | | n | n | n | n |
| 07DSGE | 64 | 64 | 67 | 19.7 | 19.7 | 20.6 | y | n | n | n | n | n |
| 08DSB | 52.5 | 57.5 | 59 | 18.1 | 19.8 | 20.4 | y | n | n | n | n | n |
| 08DSGE | 60 | 61 | 67 | 24.3 | 24.7 | 27.1 | n | n | n | n | n | y |
| 21DSB | 36.7 | 37 | 44.5 | 13.3 | 13.4 | 16.1 | n | n | n | n | n | n |
| 01DRGE | 66 | 66 | 69 | 23.9 | 23.9 | 25.0 | y | y | n | n | n | n |
| 02DRGE | 48 | 52 | 54 | 18.7 | 20.3 | 21.0 | n | n | n | n | n | n |
| 03DRGE | 54 | 58 | 71 | 17.2 | 18.5 | 22.6 | y | n | n | n | n | |
| 04DRGE | 71 | 72 | 73 | 20.5 | 20.8 | 21.1 | y | n | | n | n | n |
| 05DRGE | 60 | 62 | 65 | 19.8 | 20.4 | 21.4 | y | n | n | n | n | y |
| 06DRGE | 48 | 52 | 56 | 18.7 | 20.3 | 21.8 | n | n | n | n | n | n |
| 07DRGE | 54 | 57 | 61 | 19.5 | 20.6 | 22.1 | n | n | n | n | n | n |
| 09DRB | 43.2 | 45.5 | 44.9 | 17.9 | 18.9 | 18.6 | n | n | n | n | y | y |
| 09DRGE | | 60 | 60 | | | | n | n | n | n | n | n |
| 10DRGE | 59 | 59 | 63 | 19.9 | 19.9 | 21.2 | n | n | n | n | | n |
| 14DRGE | 60 | 62 | 66 | 20.7 | 21.4 | 22.8 | y | n | n | n | y | n |

Footnotes: BMI: body mass index.

**Supplementary Table 2. Exact p-values and test statistics for marker expression comparisons between stimulation conditions, presented in Supplementary Figure 2.**

| Population | Marker | Timepoint | Stimulation comparison | Kruskal p-value | Kruskal-Wallis Chi-Square | Degrees of freedom | Dunn's p-value | Dunn's statistic |
|---|---|---|---|---|---|---|---|---|
| $CD4^+$ | CD7 | T0 | NIL-HBHA | 0.012 | 8.75 | 2 | 0.0095 | -2.95 |
| $CD4^+$ | CD7 | T1 | NIL-HBHA | 0.029 | 7.03 | 2 | 0.031 | -2.56 |
| $CD4^+$ | CCR7 | T1 | NIL-HBHA | 0.036 | 6.62 | 2 | 0.035 | -2.51 |
| $CD8^+$ | CD7 | T0 | NIL-HBHA | 0.031 | 6.93 | 2 | 0.038 | -2.49 |
| Tgd | CD7 | T0 | NIL-HBHA | 0.036 | 6.62 | 2 | 0.042 | -2.45 |
| Tgd | CD152 | T2 | NIL-TB2 | 0.016 | 8.26 | 2 | 0.045 | -2.43 |
| Tgd | CD152 | T2 | TB2-HBHA | | | | 0.033 | 2.54 |
| DN | CD152 | T2 | TB2-HBHA | 0.030 | 6.95 | 2 | 0.030 | 2.57 |

Footnotes: here, independent, non-normal continuous variables were analyzed with the two-sided Kruskal–Wallis test with Dunn's Kruskal–Wallis Multiple Comparisons post-hoc.

**Supplementary Table 3. Exact p-values and test statistics for cluster abundance comparisons during treatment, presented in Figure 3 and Supplementary Figure 4.**

| Cluster | Timepoints | Friedman p-value | Friedman Chi-Square | Degrees of freedom | Post-hoc p-value | Post-hoc statistic |
|---|---|---|---|---|---|---|
| **NIL (n = 16 at each timepoint)** | | | | | | |
| 175 | T0-T1 | 0.012 | 8.78 | 2 | 0.008 | 16 |
| 62 | T0-T1 | 0.014 | 8.49 | 2 | 0.008 | 16 |
| 119 | T0-T1 | 0.025 | 7.35 | 2 | 0.045 | 13 |
| 171 | T0-T1 | 0.044 | 6.26 | 2 | 0.038 | 12.5 |
| 114 | T0-T1 | 0.015 | 8.4 | 2 | 0.019 | 15 |
| 98 | T0-T2 | 0.006 | 10.0 | 2 | 0.049 | 12 |
| 38 | T0-T2 | 0.004 | 11.2 | 2 | 0.021 | 14.5 |
| 176 | T0-T2 | 0.028 | 7.12 | 2 | 0.023 | 14 |
| 82 | T0-T2 | 0.033 | 6.82 | 2 | 0.031 | 14 |
| 48 | T0-T2 | 0.015 | 8.37 | 2 | 0.032 | 13.5 |
| 102 | T0-T2 | 0.003 | 11.6 | 2 | 0.019 | 14.5 |
| **TB2 (n = 18 at each timepoint)** | | | | | | |
| 164 | T0-T1 | 0.015 | 8.41 | 2 | 0.024 | 14 |
| 190 | T0-T1 | 0.002 | 12.5 | 2 | 0.013 | 16 |
| 16 | T0-T1 | 0.055 | 5.81 | 2 | 0.046 | 13.5 |
| 38 | T0-T2 | 0.007 | 9.94 | 2 | 0.025 | 15 |
| 70 | T0-T2 | 0.001 | 13.7 | 2 | 0.016 | 14.5 |
| 28 | T0-T2 | 0.013 | 8.68 | 2 | 0.012 | 16.5 |
| 37 | T0-T2 | 0.007 | 9.81 | 2 | 0.009 | 16.5 |
| 137 | T0-T2 | 0.001 | 14.9 | 2 | 0.005 | 18 |
| 74 | T0-T2 | 0.012 | 8.82 | 2 | 0.015 | 16.5 |
| 102 | T0-T2 | 0.022 | 7.61 | 2 | 0.024 | 15.5 |
| 77 | T0-T2 | 0.014 | 8.57 | 2 | 0.015 | 16 |
| **HBHA (n = 14 at each timepoint)** | | | | | | |
| 56 | T0-T1 | 0.023 | 7.53 | 2 | 0.016 | 14 |
| 27 | T0-T1 | 0.046 | 6.15 | 2 | 0.04 | 12.5 |
| 128 | T0-T1 | 0.013 | 8.74 | 2 | 0.014 | 14 |
| 69 | T0-T2 | 0.041 | 6.37 | 2 | 0.03 | 13 |
| 26 | T0-T2 | 0.02 | 7.84 | 2 | 0.018 | 14 |
| 38 | T0-T2 | 0.026 | 7.28 | 2 | 0.045 | 12.5 |
| 54 | T0-T2 | 0.03 | 7.03 | 2 | 0.019 | 13.5 |
| 172 | T0-T2 | 0.044 | 6.26 | 2 | 0.038 | 12.5 |
| 91 | T0-T2 | 0.018 | 8.04 | 2 | 0.027 | 12.5 |
| 154 | T0-T2 | 0.006 | 10.1 | 2 | 0.002 | 16.5 |
| 50 | T0-T2 | 0.039 | 6.49 | 2 | 0.032 | 13 |
| 94 | T0-T2 | 0.001 | 14.3 | 2 | 0,00 | 19 |
| 49 | T0-T2 | 0.008 | 9.69 | 2 | 0.008 | 15 |
| 160 | T0-T2 | 0.041 | 6.37 | 2 | 0.034 | 13 |
| 65 | T0-T2 | 0.039 | 6.50 | 2 | 0.04 | 12 |

Footnotes**:** For pairwise comparisons between non-independent observations at T0, T1, and T2: the two-sided Friedman rank sum test was performed followed by the Wilcoxon-Nemenyi-Thompson post-hoc.Only clusters within which significant differences were detected were represented.

**Supplementary Table 4. Pearson's correlation effect sizes (*r*) presented in Figure 5 for increased clusters.**

| | Clus 98 | Clus 38 | Clus 176 | Clus 37 | Clus 28 | Clus 70 | Clus 69 | Clus 26 | Clus 172 | Clus 91 | Clus 54 |
|---|---|---|---|---|---|---|---|---|---|---|---|
| **Clus 98** | 1 | 0.228 | -0.203 | 0.643 | 0.045 | 0.803 | 0.353 | -0.102 | -0.239 | -0.132 | 0.032 |
| **Clus 38** | 0.228 | 1 | 0.042 | 0.364 | 0.282 | 0.263 | 0.356 | -0.043 | -0.152 | 0.224 | 0.572 |
| **Clus 176** | -0.203 | 0.042 | 1 | -0.042 | -0.049 | -0.128 | -0.131 | -0.065 | -0.116 | 0.018 | -0.215 |
| **Clus 37** | 0.643 | 0.364 | -0.042 | 1 | 0.261 | 0.444 | 0.114 | -0.245 | -0.183 | 0.135 | -0.056 |
| **Clus 28** | 0.045 | 0.282 | -0.049 | 0.261 | 1 | 0.214 | 0.184 | 0.483 | 0.374 | 0.367 | 0.39 |
| **Clus 70** | 0.803 | 0.263 | -0.128 | 0.444 | 0.214 | 1 | 0.691 | 0.207 | -0.259 | 0.048 | 0.279 |
| **Clus 69** | 0.353 | 0.356 | -0.131 | 0.114 | 0.184 | 0.691 | 1 | 0.561 | -0.154 | 0.068 | 0.647 |
| **Clus 26** | -0.102 | -0.043 | -0.065 | -0.245 | 0.483 | 0.207 | 0.561 | 1 | 0.543 | 0.157 | 0.471 |
| **Clus 172** | -0.239 | -0.152 | -0.116 | -0.183 | 0.374 | -0.259 | -0.154 | 0.543 | 1 | -0.081 | 0.039 |
| **Clus 91** | -0.132 | 0.224 | 0.018 | 0.135 | 0.367 | 0.048 | 0.068 | 0.157 | -0.081 | 1 | 0.155 |
| **Clus 54** | 0.032 | 0.572 | -0.215 | -0.056 | 0.39 | 0.279 | 0.647 | 0.471 | 0.039 | 0.155 | 1 |

Footnotes: Values indicate Pearson's *r*. Correlations were calculated based on each cluster's abundance (percent of total $CD3^+$) in samples from all stimulation conditions at treatment initiation (T0). Clus: clusters. Clusters in bold indicate the clusters that were grouped together in Figure 5 for manual analysis. The associated *r* values are highlighted (orange: subgroup C corresponding to clusters, 37, 38, 70, 98; green: subgroup D corresponding to clusters 28, 54, 69).

**Supplementary Table 5. Pearson's correlation effect sizes (*r*) presented in Figure 5 for decreased clusters.**

| | **Clus 48** | **Clus 82** | **Clus 102** | **Clus 137** | **Clus 77** | **Clus 74** | **Clus 154** | **Clus 50** | **Clus 94** | **Clus 49** | **Clus 160** | **Clus 65** |
|---|---|---|---|---|---|---|---|---|---|---|---|---|
| **Clus 48** | 1 | 0.377 | 0.412 | 0.137 | 0.462 | 0.725 | 0.485 | -0.093 | 0.711 | 0.565 | 0.286 | 0.372 |
| **Clus 82** | 0.377 | 1 | -0.006 | 0.189 | 0.064 | 0.48 | 0.456 | 0.318 | 0.615 | 0.424 | 0.114 | 0.686 |
| **Clus 102** | 0.412 | -0.006 | 1 | -0.062 | 0.635 | 0.485 | 0.093 | -0.365 | 0.206 | -0.027 | 0.585 | -0.087 |
| **Clus 137** | 0.137 | 0.189 | -0.062 | 1 | -0.072 | -0.054 | 0.226 | -0.113 | 0.066 | 0.237 | -0.008 | -0.076 |
| **Clus 77** | 0.462 | 0.064 | 0.635 | -0.072 | 1 | 0.614 | 0.189 | -0.239 | 0.352 | 0.076 | 0.6 | -0.027 |
| **Clus 74** | 0.725 | 0.48 | 0.485 | -0.054 | 0.614 | 1 | 0.261 | -0.048 | 0.678 | 0.311 | 0.318 | 0.375 |
| **Clus 154** | 0.485 | 0.456 | 0.093 | 0.226 | 0.189 | 0.261 | 1 | 0.288 | 0.471 | 0.513 | 0.145 | 0.64 |
| **Clus 50** | -0.093 | 0.318 | -0.365 | -0.113 | -0.239 | -0.048 | 0.288 | 1 | 0.124 | 0.263 | -0.199 | 0.673 |
| **Clus 94** | 0.711 | 0.615 | 0.206 | 0.066 | 0.352 | 0.678 | 0.471 | 0.124 | 1 | 0.324 | 0.241 | 0.637 |
| **Clus 49** | 0.565 | 0.424 | -0.027 | 0.237 | 0.076 | 0.311 | 0.513 | 0.263 | 0.324 | 1 | -0.063 | 0.452 |
| **Clus 160** | 0.286 | 0.114 | 0.585 | -0.008 | 0.6 | 0.318 | 0.145 | -0.199 | 0.241 | -0.063 | 1 | -0.017 |
| **Clus 65** | 0.372 | 0.686 | -0.087 | -0.076 | -0.027 | 0.375 | 0.64 | 0.673 | 0.637 | 0.452 | -0.017 | 1 |

Footnotes: Values indicate Pearson's *r*. Correlations were calculated based on each cluster's abundance (percent of total $CD3^+$) in samples from all stimulation conditions at treatment initiation (T0). Clus: clusters. Clusters in bold indicate the clusters that were grouped together in Figure 5 for manual analysis. The associated *r* values are highlighted (green: subgroup A corresponding to clusters, 49, 50, 65, and 154; blue: subgroup B corresponding to clusters 74, 102, 160).

**Supplementary Table 6. Exact p-values and test statistics for cluster abundance comparisons between fast and slow converters, presented in Figure 6 and Supplementary Figure 5.**

| Cluster | Timepoint | Stimulation | N(fast converters) | N(slow converters) | U statistic | p-value |
|---|---|---|---|---|---|---|
| **T0 - significance threshold set at p < 0.026** | | | | | | |
| 117 | T0 | NIL | 12 | 4 | 45.5 | 0.0094 |
| 132 | T0 | NIL | 12 | 4 | 46 | 0.0079 |
| 134 | T0 | NIL | 12 | 4 | 45.5 | 0.0099 |
| 147 | T0 | NIL | 12 | 4 | 43 | 0.023 |
| 148 | T0 | NIL | 12 | 4 | 43.5 | 0.019 |
| 186 | T0 | NIL | 12 | 4 | 3.5 | 0.0089 |
| 48 | T0 | NIL | 12 | 4 | 44 | 0.017 |
| 62 | T0 | NIL | 12 | 4 | 46 | 0.0087 |
| 76 | T0 | NIL | 12 | 4 | 43.5 | 0.020 |
| 88 | T0 | NIL | 12 | 4 | 43.5 | 0.020 |
| 105 | T0 | TB2 | 14 | 4 | 54 | 0.0061 |
| 118 | T0 | TB2 | 14 | 4 | 51.5 | 0.012 |
| 134 | T0 | TB2 | 14 | 4 | 52.5 | 0.010 |
| 147 | T0 | TB2 | 14 | 4 | 50 | 0.021 |
| 148 | T0 | TB2 | 14 | 4 | 49.5 | 0.025 |
| 32 | T0 | TB2 | 14 | 4 | 50 | 0.020 |
| 74 | T0 | TB2 | 14 | 4 | 49.5 | 0.025 |
| 75 | T0 | TB2 | 14 | 4 | 54.5 | 0.0053 |
| 89 | T0 | TB2 | 14 | 4 | 50 | 0.020 |
| 90 | T0 | TB2 | 14 | 4 | 52 | 0.010 |
| 102 | T0 | HBHA | 11 | 3 | 32 | 0.018 |
| 132 | T0 | HBHA | 11 | 3 | 31.5 | 0.021 |
| 134 | T0 | HBHA | 11 | 3 | 31.5 | 0.023 |
| 148 | T0 | HBHA | 11 | 3 | 33 | 0.012 |
| 182 | T0 | HBHA | 11 | 3 | 2 | 0.015 |
| 183 | T0 | HBHA | 11 | 3 | 0 | 0.010 |
| 52 | T0 | HBHA | 11 | 3 | 31.5 | 0.023 |
| 62 | T0 | HBHA | 11 | 3 | 32.5 | 0.015 |
| 75 | T0 | HBHA | 11 | 3 | 31.5 | 0.021 |

**Supplementary Table 6 - continued**

| T1 - significance threshold set at p < 0.031 | | | | | | |
|---|---|---|---|---|---|---|
| 116 | T1 | NIL | 12 | 4 | 43 | 0.023 |
| 120 | T1 | NIL | 12 | 4 | 43.5 | 0.019 |
| 180 | T1 | NIL | 12 | 4 | 6 | 0.023 |
| 75 | T1 | NIL | 12 | 4 | 43.5 | 0.020 |
| 11 | T1 | TB2 | 14 | 4 | 7.5 | 0.031 |
| 117 | T1 | TB2 | 14 | 4 | 50 | 0.021 |
| 119 | T1 | TB2 | 14 | 4 | 49 | 0.027 |
| 125 | T1 | TB2 | 14 | 4 | 5.5 | 0.018 |
| 132 | T1 | TB2 | 14 | 4 | 49.5 | 0.025 |
| 134 | T1 | TB2 | 14 | 4 | 55 | 0.0047 |
| 146 | T1 | TB2 | 14 | 4 | 49.5 | 0.025 |
| 147 | T1 | TB2 | 14 | 4 | 53.5 | 0.0075 |
| 148 | T1 | TB2 | 14 | 4 | 52.5 | 0.010 |
| 166 | T1 | TB2 | 14 | 4 | 7.5 | 0.019 |
| 171 | T1 | TB2 | 14 | 4 | 6 | 0.021 |
| 178 | T1 | TB2 | 14 | 4 | 6 | 0.020 |
| 180 | T1 | TB2 | 14 | 4 | 7.5 | 0.028 |
| 4 | T1 | TB2 | 14 | 4 | 6 | 0.021 |
| 57 | T1 | TB2 | 14 | 4 | 5 | 0.016 |
| 62 | T1 | TB2 | 14 | 4 | 51 | 0.016 |
| 64 | T1 | TB2 | 14 | 4 | 49.5 | 0.025 |
| 76 | T1 | TB2 | 14 | 4 | 51 | 0.016 |
| 89 | T1 | TB2 | 14 | 4 | 49.5 | 0.023 |
| 117 | T1 | HBHA | 11 | 3 | 33 | 0.011 |
| 12 | T1 | HBHA | 11 | 3 | 1.5 | 0.022 |
| 147 | T1 | HBHA | 11 | 3 | 31.5 | 0.021 |
| 180 | T1 | HBHA | 11 | 3 | 1.5 | 0.020 |
| 34 | T1 | HBHA | 11 | 3 | 31 | 0.028 |
| 62 | T1 | HBHA | 11 | 3 | 32 | 0.018 |

**Supplementary Table 6 - continued**

| T2 - significance threshold set at p < 0.013 | | | | | | |
|---|---|---|---|---|---|---|
| 112 | T2 | NIL | 12 | 4 | 2 | 0.0073 |
| 12 | T2 | NIL | 12 | 4 | 2 | 0.0052 |
| 134 | T2 | NIL | 12 | 4 | 46.5 | 0.0074 |
| 14 | T2 | NIL | 12 | 4 | 2 | 0.0073 |
| 169 | T2 | NIL | 12 | 4 | 0 | 0.0031 |
| 181 | T2 | NIL | 12 | 4 | 3 | 0.012 |
| 182 | T2 | NIL | 12 | 4 | 0.5 | 0.0021 |
| 183 | T2 | NIL | 12 | 4 | 1 | 0.0049 |
| 60 | T2 | NIL | 12 | 4 | 45 | 0.012 |
| 62 | T2 | NIL | 12 | 4 | 47 | 0.0063 |
| 66 | T2 | NIL | 12 | 4 | 45.5 | 0.010 |
| 67 | T2 | NIL | 12 | 4 | 45 | 0.012 |
| 74 | T2 | NIL | 12 | 4 | 47 | 0.0059 |
| 81 | T2 | NIL | 12 | 4 | 45 | 0.012 |
| 112 | T2 | TB2 | 14 | 4 | 4 | 0.012 |
| 12 | T2 | TB2 | 14 | 4 | 3 | 0.0085 |
| 134 | T2 | TB2 | 14 | 4 | 52 | 0.012 |
| 148 | T2 | TB2 | 14 | 4 | 53 | 0.0088 |
| 166 | T2 | TB2 | 14 | 4 | 2 | 0.0045 |
| 169 | T2 | TB2 | 14 | 4 | 4 | 0.011 |
| 180 | T2 | TB2 | 14 | 4 | 2 | 0.0062 |
| 181 | T2 | TB2 | 14 | 4 | 1.5 | 0.0055 |
| 182 | T2 | TB2 | 14 | 4 | 1 | 0.0035 |
| 183 | T2 | TB2 | 14 | 4 | 3 | 0.0064 |
| 62 | T2 | TB2 | 14 | 4 | 55 | 0.0047 |
| 88 | T2 | TB2 | 14 | 4 | 52 | 0.012 |
| 94 | T2 | TB2 | 14 | 4 | 54 | 0.0059 |
| 98 | T2 | TB2 | 14 | 4 | 3.5 | 0.010 |
| 183 | T2 | HBHA | 11 | 3 | 0 | 0.011 |

Footnotes**:** For comparisons between non-normal, independent continuous variables at T0, T1, and T2 separately, the two-sided Mann-Whitney U test was performed. For discovery of clusters with significantly different abundance, conservative corrections for multiple comparisons (*e.g.* Benjamini-Hochberg) were not used in order to minimize type II errors. Instead, all p-values were computed for each timepoint, and the p-value corresponding to the null hypothesis being rejected in 5% of all comparisons was used as the significance threshold instead of 0.05. This novel significance threshold enabled to control type I error while maintaining an exploratory approach. Only clusters within which significant differences were detected were represented.

**Supplementary Table 7. Exact p-values and test statistics for comparison of PCA scores between fast and slow converters, presented in Figure 7 and Supplementary Figure 7.**

| Timepoint | N(fast converters) | N(slow converters) | U statistic | p-value |
|---|---|---|---|---|
| **Figure 7 - within selected clusters** | | | | |
| **Component 1** | | | | |
| T0 | 25 | 7 | 34 | 0.013 |
| T1 | 25 | 7 | 28 | 0.0051 |
| T2 | 25 | 7 | 25 | 0.0029 |
| **Component 2** | | | | |
| T0 | 25 | 7 | 130 | 0.0542 |
| T1 | 25 | 7 | 72 | 0.503 |
| T2 | 25 | 7 | 146 | 0.00602 |
| **Supp. Figure 7 - within all $CD3^+$ events** | | | | |
| **Component 1** | | | | |
| T0 | 25 | 7 | 24 | 0.00247 |
| T1 | 25 | 7 | 30 | 0.00709 |
| T2 | 25 | 7 | 22 | 0.00166 |
| **Component 2** | | | | |
| T0 | 25 | 7 | 70 | 0.447 |
| T1 | 25 | 7 | 101 | 0.562 |
| T2 | 25 | 7 | 58 | 0.191 |

Footnotes: For comparisons between non-normal, independent continuous variables at T0, T1, and T2 separately, the two-sided Mann-Whitney U test was performed.

**Supplementary Table 8. Mass cytometry panel components.**

Panel will be accessible upon author request. All antibodies were supplied by Fluidigm.

**Supplementary Table 9. Definition of clustering channels and expected cell subpopulations for dimension reduction and automated clustering of $CD3^{+}$ T-cells.**

| Clustering channels | Expected cell subpopulations | Phenotype |
|---|---|---|
| CD4, CD8, CCR7, CD45RA, CD161, CD26, TCRgd, TCRVa7.2, CD25 | Gamma delta T-cells | $CD3^{+}$ $TCR\gamma\delta^{+}$ |
| | MAIT-cells | $CD3^{+}$ $CD4^{-}$ $CD8^{+}$ $CD26^{+}$ $CD161^{+}$ $TCRV\alpha7.2^{+}$ |
| | Naive $CD8^{+}$ T-cells | $CD3^{+}$ $CD4^{-}$ $CD8^{+}$ $CCR7^{+}$ $CD45RA^{+}$ |
| | Effector memory $CD8^{+}$ T-cells | $CD3^{+}$ $CD4^{-}$ $CD8^{+}$ $CCR7^{-}$ $CD45RA^{-}$ |
| | Central memory $CD8^{+}$ T-cells | $CD3^{+}$ $CD4^{-}$ $CD8^{+}$ $CCR7^{+}$ $CD45RA^{-}$ |
| | TEMRA $CD8^{+}$ T-cells | $CD3^{+}$ $CD4^{-}$ $CD8^{+}$ $CCR7^{-}$ $CD45RA^{+}$ |
| | Naive $CD4^{+}$ T-cells | $CD3^{+}$ $CD4^{+}$ $CD8^{-}$ $CCR7^{+}CD45RA^{+}$ |
| | Effector memory $CD4^{+}$ T-cells | $CD3^{+}$ $CD4^{+}$ $CD8^{-}$ $CCR7^{-}CD45RA^{-}$ |
| | Central memory $CD4^{+}$ T-cells | $CD3^{+}$ $CD4^{+}$ $CD8^{-}$ $CCR7^{+}CD45RA^{-}$ |
| | TEMRA $CD4^{+}$ T-cells | $CD3^{+}$ $CD4^{+}$ $CD8^{-}$ $CCR7^{-}$ $CD45RA^{+}$ |
| | Treg | $CD3^{+}$ $CD4^{+}$ $CD25^{+}$ $IL7Ra^{-}$ |
| | Double negative T-cells | $CD3^{+}$ $CD4^{-}$ $CD8^{-}$ |

Footnotes: only lineage-defining markers are presented in this table. MAIT: mucosal-associated invariant T-cells. TEMRA: terminally differentiated effectors re-expressing CD45RA. NK: natural killer cells. NKT: natural killer T-cells. Treg: T regulators.
